# Fine-tuning m6A and METTL3 levels have profound impact on cellular proliferation and protein synthesis

**DOI:** 10.64898/2026.04.14.718245

**Authors:** Samir Watson, Johanna Luige, Sarah N’Da Konan, Peter Alling Dam, Alessandro Brambilla, Helene Overvad Pedersen, Ulf Andersson Vang Ørom

**Author notes:** Corresponding authors (SW), (UAVØ). Further information and requests for resources and reagents should be directed to and will be fulfilled by the Lead Contact (Ulf Andersson Vang Ørom –).

## Abstract

The RNA modification m6A is the most abundant internal RNA modification in eukaryotic mRNAs and long non-coding RNAs and has been implicated in diverse and important biological processes. Notably, m6A has been associated with both pro and anti-tumorigenic roles depending on cellular and biological context. In basal-like triple negative breast cancer (TNBC), heterozygous loss of the m6A methyltransferase METTL3 and increased levels of the m6A demethylase FTO are associated with poor prognosis and an increased risk of metastasis. Here, using CRISPR generated METTL3 heterozygous knockout TNBC cell lines (MDA-MB-468) and Nanopore direct RNA sequencing, we characterise transcriptome-wide changes in m6A modification patterns following partial loss of METTL3. We reveal that partial loss of global m6A is associated with preferential changes in the methylation status of transcripts involved in translational control, leading to an increase in translational output and proliferative capacity. In contrast, strong pharmacologic inhibition of METTL3 suppresses translation and proliferation. Our findings highlight how m6A levels differentially regulate gene expression in a dose dependent manner and provides a deeper molecular understanding of the RNA modification m6A and its role in fine-tuning translation and affecting both tumorigenesis and cancer progression.

## Introduction

N6-methyladenosine (m6A) is the most abundant modification found in mRNAs with an abundance of between 0.1-0.4% of all adenosines and between 25-60% of all transcripts are found to contain m6A modified nucleotides^1–4^. m6A is deposited by the methyltransferase complex, a large multi-subunit protein complex consisting of three core proteins, METTL3, METTL14 and WTAP and several accessory proteins, including RBM15, ZC3H13, and VIRMA ^5^. METTL3 is the enzyme responsible for carrying out the methylation of adenosines, often occurring at a DRACH (IUPAC degenerate nucleotide code) consensus motif ^6^. m6A deposition is a dynamic process as it can be removed by the m6A eraser proteins FTO and ALKBH5 ^7,8^. In addition, m6A is recognised by a diverse set of reader proteins including the YTH family proteins, HNRNPC/G, HNRNPA2B1, IGF2BP1-3 amongst others, that bind to m6A containing RNAs leading to functional outcomes on the bound RNA, including increase in translation and RNA degradation depending on both the reader and the m6A context^9–15^.

Previous studies have primarily focused on acute or complete knockouts/knockdowns of METTL3 to study the effects of m6A on RNA fate. However, owing to the revelation that METTL3 is embryonic lethal in mouse and is an essential gene in most human cell lines, generation of acute or complete METTL3 KO/KD lines is often not possible, and in most cases not physiologically relevant ^16–19^. Several studies generating CRISPR/Cas9 METTL3 knockouts in mouse and in human cell lines have reported a wide range of residual m6A on mRNAs that were originally attributed to other methyltransferases compensating for loss of m6A^20–23^. However, a comprehensive study of this phenomenon revealed that many apparent CRISPR knockout cell lines rely on alternative splice METTL3 isoforms to maintain viability ^19^. The consequences of partial loss of METTL3 activity have not been systematically followed although several studies on cancer have revealed that dysregulation of the m6A RNA modification on the transcriptome has a bearing on negative outcomes for patients^24–26^. As such METTL3, a key component of the m6A methyltransferase complex, has been implicated as both a tumour suppressor and an oncogene in different cancers ^24,27^.

Triple negative breast cancer (TNBC) lacks estrogen, progesterone and human epidermal growth factor 2 receptors and is the most aggressive subtype of breast cancer causing 15-20% of all breast cancers with the poorest prognosis due to lack of molecular targeted treatment^28–30^. It is highly associated with overweight and obesity – which is also true for other breast cancers^31–33^. TNBCs are difficult to treat and highly invasive with a high recurrence rate. In TNBC, reduced expression of METTL3 coincides with an increase in metastasis and breast cancer progression ^27,34^. However, other studies have revealed that METTL3 is overexpressed in other breast cancers ^35^. The conflicting evidence points to a complex interplay between m6A and cancer.

It has been proposed that m6A modification levels are important as it has been linked to numerous pathologies including cancers, neurodegenerative diseases and autoimmune disorders^36–38^. Moreover, both the position of m6A marks along the mRNA and their timing of deposition appear to influence their regulatory effects^39^. Thus, ample evidence supports a complex m6A landscape, which can drastically alter the fates of target mRNAs – the disruption of which may contribute to disease development.

Here, we investigate how partial loss of METTL3 alters the m6A landscape and downstream growth signalling in TNBC. Using CRISPR-engineered METTL3 heterozygous knockout MDA-MB-468 cells together with pharmacological inhibition, we combine Nanopore direct RNA sequencing with functional analyses of translation and proliferation. We show that modest reductions in global m6A levels preferentially affect a subset of transcripts linked to translation and cell growth, coinciding with increased translational output and proliferative capacity, whereas stronger suppression of METTL3 activity impairs protein synthesis and cell growth. These findings reveal a dose-dependent role for m6A in tuning translational control and highlight how partial disruption of METTL3 can engage growth-promoting pathways in cancer cells, providing mechanistic insight into the context-dependent roles of m6A in tumour progression.

## Results

### Partial loss of METTL3 increases cell proliferation in MDA-MB-468 cells

To elucidate the effect of an altered m6A equilibrium, we generated heterozygous knockout cell lines using the CRISPR CAS9 system, targeting exon 1 of METTL3, in a TNBC cell line, MDA-MB-468. These cells are more fibroblast-like, less invasive and less malignant than other frequently used breast cancer cell lines such as MDA-MB-231 ^40,41^. In the heterozygous knockout cell line, HKO (Figure 1a), one allele exhibits a deletion of 317 bp, encompassing the entire METTL3 exon 1. While the other allele has a 67 bp deletion located upstream of exon 1, which does not alter the coding sequence (Figure 1b, c). Q-RT-PCR of METTL3 reveals a 69% reduction in METTL3 mRNA levels whilst Western blotting reveals a 65% reduction in METTL3 protein levels (Figure 1d-f). Treatment using the METTL3-specific chemical inhibitor, STM2457, at a concentration of 0.145 µM, which corresponds to the level of m6A depletion in the HKO cell lines, did not lead to any changes in METTL3 protein or RNA levels (Figure 1, d, e, Supplementary Figure 1a). RNA mass spectrometry of mRNA shows that despite a robust reduction in protein levels, there is a modest 22.5% reduction in m6A levels (Figure 1g), suggesting a methyltransferase system that is relatively robust against fluctuations in METTL3 levels. Western blot analysis of the m6A writer machinery reveals no changes to the expression of RBM15 and VIRMA (Figure 1e). However, we observe a marked decrease in both METTL14 steady state RNA and protein levels (Figure 1 d, e). The Western blots are also carried out on protein lysates from cells treated with the specific METTL3 inhibitor STM2457 showing that the reduction in METTL14 levels is independent of m6A levels and suggesting that METTL14 protein stability depends on METTL3 protein levels, rather than on its activity (Figure 1e), as has also been reported previously^42–44^. Additionally, we investigated the mRNA levels of the m6A erasers FTO and ALKBH5 as well as the MTC component WTAP and the methionine adenosyltransferase, MAT2A, which has been mechanistically shown to be regulated by mTORC1, and is essential for maintenance of cellular SAM levels required by METTL3 for RNA methylation^45^. We do not observe any significant changes to steady state RNA levels for these genes using the METTL3 inhibitor or in our HKO lines (Supplementary Figure 1 b)

**Figure 1.**
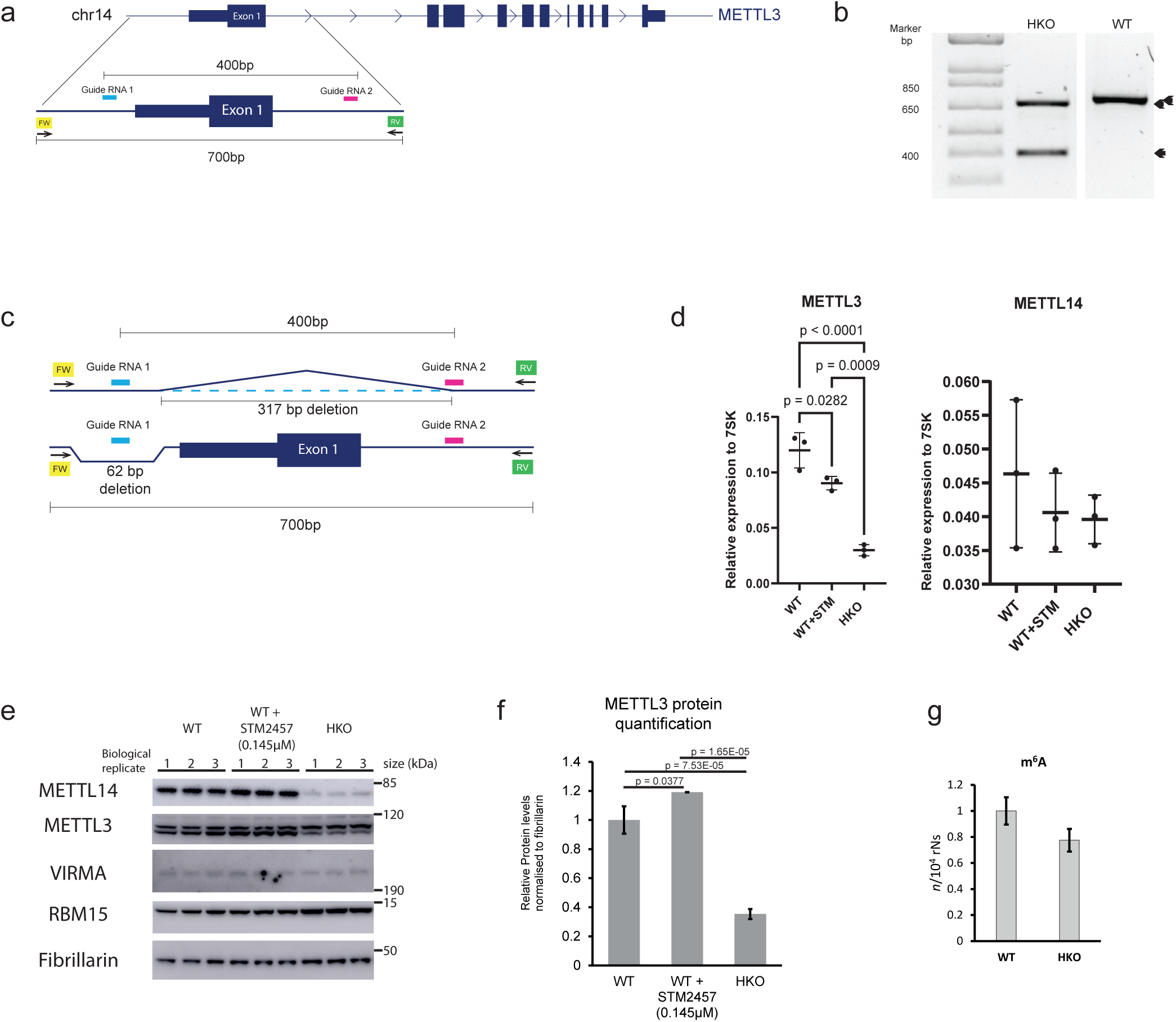
Generation of heterozygous knockout of METTL3 in MDA-MB-468 TNBC cells. **a)** Design and location of CRISPR guide RNAs (pink and blue) and genotyping primers (yellow and green) targeting exon 1 of METTL3 **b)** Agarose gel showing the genotyping PCR products amplified from the CRISPR-Cas9 targeted region of a heterozygous knockout clone (HKO) and the parental cell line MDA-MB-468 (WT). **c)** Representation of METTL3 exon 1 of HKO showing the representative locations of CRISPR heterozygous deletions. **d)** qPCR analysis of METTL3 and METTL14 gene expression in WT and METTL3 HKO cells. P-value by one-way ANOVA followed by Tukey HSD post hoc multiple comparison test. **e)** Western blot analysis of METTL3, METTL14, VIRMA, RBM15 protein expression wild type and METTL3 +/− cells. Fibrillarin serves as a loading control. **f)** Western blot quantification of METTL3 on WT treated with 0.145µM STM2457 (n=3) or METTL3 HKO lines (n=3), compared to WT (n=3). Protein levels are represented as mean ratio values quantified from protein bands of METTL3 (lower band) versus Fibrillarin compared to WT. P-values calculated by one-way ANOVA followed by Tukey HSD post hoc multiple comparison test. **g)** m6A/A ratio of poly(A) purified mRNA as determined by LC–MS/MS in KO (n=2) and WT(n=2) cells.

### METTL3 dosage alters proliferation capacity in MDA-MB-468 cells

We examined how altering METTL3 dosage or catalytic activity affects proliferation capacity in MDA-MB-468 TNBC cells by comparing METTL3 heterozygous knockout (HKO) cells to WT controls and by pharmacologic inhibition using STM2457 (0.145 µM or 3 µM) or the weak analogue STM2120 (3 µM), which is approximately 1000-fold less potent than STM2457. Cells were pre-treated for 24 h and then exposed to mitogenic, metabolic, or inhibitory co-treatments (EGF, EGF plus dexamethasone, insulin, rapamycin, metformin, or control) across four glucose concentrations (0, 5.5, 15, and 25 mM). Proliferation capacity was quantified using longitudinal CellTiter-Glo luminescence measurements collected on separate plates from 24 to 120 h and summarized primarily as absolute area under the curve (AUC; 24–120 h) (Fig. 2A).

**Figure 2.**
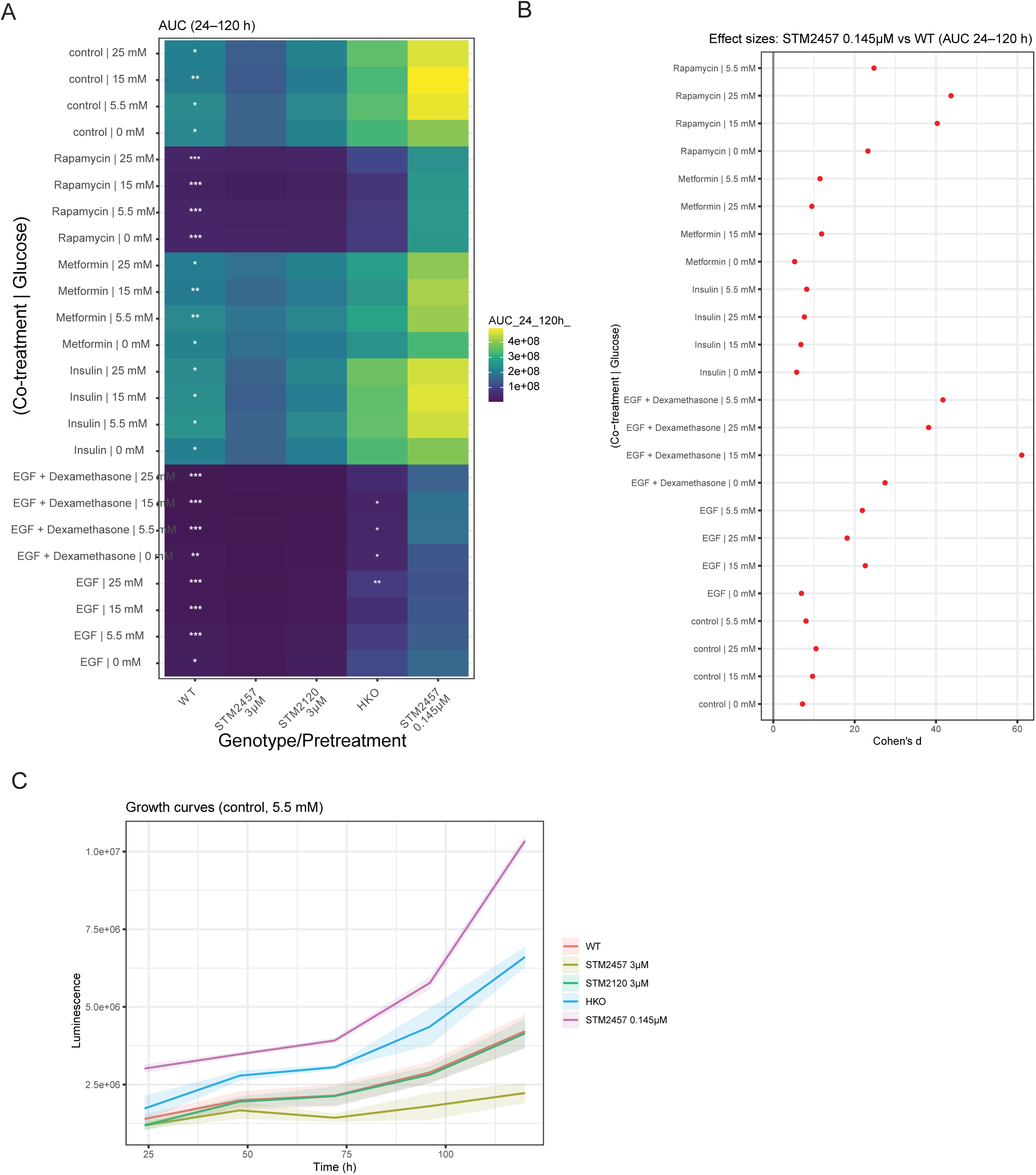
METTL3 perturbation alters proliferation capacity in MDA-MB-468 TNBC cells. **a)** Heatmap showing absolute area under the curve (AUC; 24–120 h) across co-treatment and glucose conditions (rows) and genotype or pretreatment (columns: WT, STM2457 3 µM, STM2120 3 µM, METTL3 heterozygous knockout [HKO], STM2457 0.145 µM). Asterisks indicate BH-FDR–adjusted significance relative to WT (*** ≤ 0.001; ** ≤ 0.01; * ≤ 0.05). **b)** Effect sizes (Cohen’s d) comparing STM2457 0.145 µM to WT across all conditions (red denotes FDR ≤ 0.05).**c)** Absolute growth curves (mean ± SEM) for MDA-MB-468 WT cells treated with STM2457 (3 µM or 0.145 µM), STM2120 (3 µM), or METTL3 HKO cells, cultured in 5.5 mM glucose. Together, panels (A–C) demonstrate that METTL3 HKO increases growth capacity relative to WT under mitogenic and nutrient-rich conditions, STM2457 3 µM suppresses growth capacity in STM-only and selected co-treatment contexts, and STM2457 0.145 µM produces modest effects relative **to** WT.

Across conditions, METTL3 HKO cells generally exhibited higher absolute AUC than WT cells (Fig. 2A), with the largest differences observed under strong mitogenic stimulation (EGF ± dexamethasone) and at elevated glucose concentrations (15–25 mM). These patterns indicate that mitogenic signalling and nutrient abundance amplify the increased proliferation capacity associated with partial METTL3 loss. In contrast, pharmacologic inhibition with STM2457 at 3 µM reduced AUC relative to WT in STM-only and selected co-treatment contexts (Fig. 2A), consistent with suppression of proliferation capacity under strong catalytic inhibition. STM2457 at 0.145 µM produced minimal to modest changes in AUC relative to WT across most conditions, while STM2120 at 3 µM closely tracked WT behaviour, consistent with its substantially lower inhibitory potency (Fig. 2A). Notably, STM2457 at 0.145 µM induces a level of global m6A depletion comparable to that observed in METTL3 HKO cells, indicating that this dose represents a catalytic-equivalent perturbation rather than maximal METTL3 inhibition.

Representative luminescence trajectories at 5.5 mM glucose further illustrate these trends (Fig. 2B). HKO cells displayed increased signal accumulation over time relative to WT, whereas STM2457 at 3 µM suppressed signal accumulation, and STM2457 at 0.145 µM and STM2120 at 3 µM exhibited trajectories similar to WT.

Rapamycin broadly reduced proliferation capacity across genotypes and treatment conditions (Fig. 2A); however, HKO cells retained a relative advantage over WT, suggesting that the METTL3-dependent phenotype is not solely mediated through mTORC1 signalling. Insulin and metformin each reduced or modulated AUC in a manner consistent with activation of PI3K/AKT and AMPK signalling, respectively, yet HKO cells frequently remained above WT under these conditions (Fig. 2A), supporting a robust increase in proliferation capacity associated with partial METTL3 loss across diverse signalling contexts.

Effect size analysis (Cohen’s d) highlighted a practically meaningful shift for STM2457 at 0.145 µM relative to WT across multiple conditions, with statistically significant contexts following false discovery rate correction indicated in Fig. 2B. Stronger negative effect sizes were observed for STM2457 at 3 µM in selected contexts, consistent with pronounced suppression of proliferation capacity at higher inhibitor concentration. Normalized luminescence trajectories, shown in Supplementary Fig. S3, illustrate the dynamic behaviour underlying the absolute AUC measurements while minimizing technical variability due to plate and seeding effects and support the same qualitative conclusions.

Taken together, these data show that METTL3 dosage and catalytic activity bidirectionally tune proliferation capacity in MDA-MB-468 cells. Heterozygous loss of METTL3 consistently increases absolute AUC (24–120 h) relative to WT across nutrient and signalling contexts, with the largest gains observed under EGF or EGF plus dexamethasone and at elevated glucose concentrations, where mitogenic and metabolic inputs are strongest. In contrast, pharmacologic inhibition with STM2457 reduces proliferation capacity in a dose-dependent manner: 3 µM produces clear suppression, whereas 0.145 µM—matched to HKO-like global m6A depletion—elicits modest effects that align with inhibitor potency and pathway context. Importantly, the HKO-associated advantage persists under rapamycin treatment, indicating that this phenotype is not solely mTORC1-driven, and remains evident in insulin- and metformin-treated conditions that modulate PI3K/AKT and AMPK signalling. Normalised luminescence trajectories (24 h baseline = 1; AUC 0–120 h) corroborate these conclusions while reducing technical variability due to plate and seeding effects. Overall, the results support a model in which partial METTL3 loss elevates baseline proliferation capacity, whereas strong catalytic inhibition suppresses it, with the magnitude and detectability of effects shaped by mitogenic strength and glucose availability.

To determine whether altered proliferation capacity reflected changes in cell viability rather than growth rate, we assessed apoptosis and necrosis by Annexin V/7-AAD staining following partial genetic or pharmacologic perturbation of METTL3. Neither heterozygous loss of METTL3 nor treatment with STM2457 at either 0.145 µM or 3 µM produced a consistent increase in early or late apoptotic, necrotic, or non-viable cell populations relative to WT controls (Supplementary Fig. 2). Across biological replicates, the proportion of live cells remained comparable between conditions, indicating that the proliferation phenotypes observed upon partial METTL3 loss or inhibition are not driven by drug- or genotype-induced cytotoxicity.

### Partial loss of METTL3 leads to an altered m6A landscape

To investigate how heterozygous loss of METTL3 alters m6A deposition in TNBC cells, we performed Oxford Nanopore direct RNA sequencing (DRS) on three biological replicates of WT and heterozygous knockout (HKO) cell lines. Sequencing yielded sufficient read depth, read length, and base quality across all samples to enable robust detection of m6A modifications and differential methylation analysis.

To characterise m6A changes at complementary levels, we applied three nanopore-based analysis approaches. First, we quantified global changes in canonical m6A at DRACH motifs using the m6Anet pipeline. Second, we identified reproducibly differentially modified sites across replicates using xPore and Nanocompore. Finally, we estimated methylation stoichiometry at selected high-confidence sites using a modified nanoRMS pipeline to assess the magnitude of differential modification.

Global analysis using m6Anet revealed a consistent reduction in m6A modification in HKO cells, with a 16–20% decrease in the average modification ratio across all DRACH motifs compared to WT (Figure 3a). This indicates that partial loss of METTL3 leads to a measurable but incomplete reduction in canonical m6A deposition.

**Figure 3.**
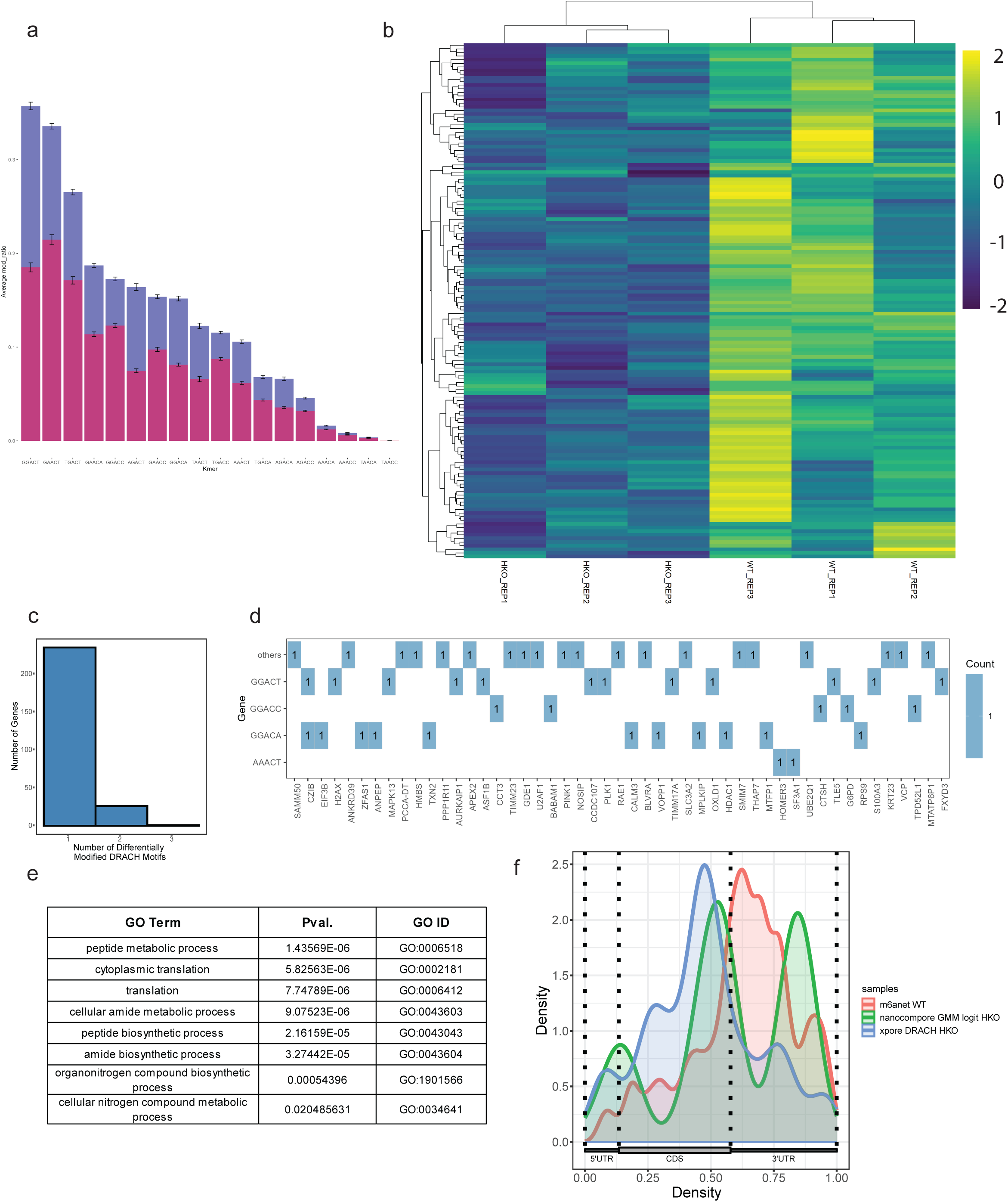
m^6^A profiling using nanopore Direct RNA sequencing. **a)** Comparison of the average modification ratio predicted by m6Anet between WT (purple) and HKO-1 and HKO-2 combined (pink) on DRACH kmers. **b)** Heatmap of significantly differentially modified DRACH kmers (from 290 sites) identified by Xpore, comparing modification rate values across genes with consistent expression across all samples (>1 reads in all samples). **c)** Frequency of differentially modified DRACH kmers per gene identified by xPore (pval <0.05, Z-score >0, assignment, lower). **d)** Frequency at which the top 5 most frequently modified DRACH kmers as identified by m6Anet appear in the 50 genes that exhibit the highest rates of differential modification. All other DRACH kmers are combined in the others row. **e)** Top enriched biological process ontologies of differentially modified genes identified by XPore (pval <0.05, Holm-Bonferroni correction). **f)** Metagene profiles of the distribution of m6A modified predicted by m6Anet sites (red, probability >0.9) and differentially modified sites identified by nanocompore using the GMM logit statistical test (green, pval <0.01, LOR>0) and xPore (blue, pval <0.05).

To identify site-specific changes, we performed differential modification analysis using xPore, which identified 290 significantly differentially methylated DRACH sites (Fig. 3b; Supplementary Table 1). Unsupervised hierarchical clustering of consistently expressed transcripts containing these sites separated WT and HKO samples, demonstrating a transcriptome-wide shift in methylation patterns upon METTL3 heterozygosity (Figure 3b). Most differentially methylated transcripts contained a single affected DRACH site (Figure 3c), including among the top-ranked genes with the largest differential modification rates (Figure 3d).

Functional enrichment analysis of transcripts containing differentially modified DRACH sites revealed pathways sensitive to partial METTL3 loss. Gene ontology analysis, as well as KEGG and Reactome pathway enrichment, showed significant overrepresentation of genes involved in translation and metabolism, a pattern observed independently using both xPore and Nanocompore outputs (Figure 3e; Supplementary Tables 1–3). Additional enrichment was observed for mitochondrial translation, cell cycle regulation, and nonsense-mediated decay pathways (Supplementary Table 3).

Metagene analysis of predicted m6A sites identified by m6Anet (probability > 0.9) showed that m6A distribution in WT cells was enriched near stop codons and within 3′ untranslated regions, consistent with previously reported profiles in HEK293T and HCT116 cells (Figure 3f, red; Supplementary Table 4). Differentially modified DRACH sites identified by xPore were enriched within gene bodies, with a peak immediately upstream of the stop codon and enrichment in the 3′UTR (Figure 3f, blue). Nanocompore-identified differential sites displayed three enrichment peaks, located near the start codon, upstream of the stop codon, and within the 3′UTR (Figure 3f, green).

To estimate the magnitude of differential m6A modification at individual sites, we applied a modified version of the nanoRMS pipeline to high-confidence DRACH sites identified by xPore and Nanocompore. As our experimental system involves relatively modest changes in total m6A levels, stoichiometry estimation relied on unsupervised KMEANS clustering of nanopore signal features, where clusters were interpreted based on relative changes in abundance between WT and HKO samples.

We first examined a differentially modified DRACH site in *Cyclin D3*, identified by xPore (p < 0.05; Supplementary Tables 1 and 2). Current intensity density at position 115 revealed a shift in signal distribution in HKO cells compared to WT, both at the central site and across a 15-nucleotide window surrounding the modification (Figure 4f,g). Principal component analysis of per-read signal features across this window showed a clear separation between WT and HKO reads, indicating systematic differences in nanopore signal associated with reduced m6A levels (Figure 4h). KMEANS clustering estimated a 26.7% reduction in m6A stoichiometry in HKO cells relative to WT (Figure 4i,j), consistent with the 21.7% differential modification rate estimated independently by xPore.

**Figure 4.**
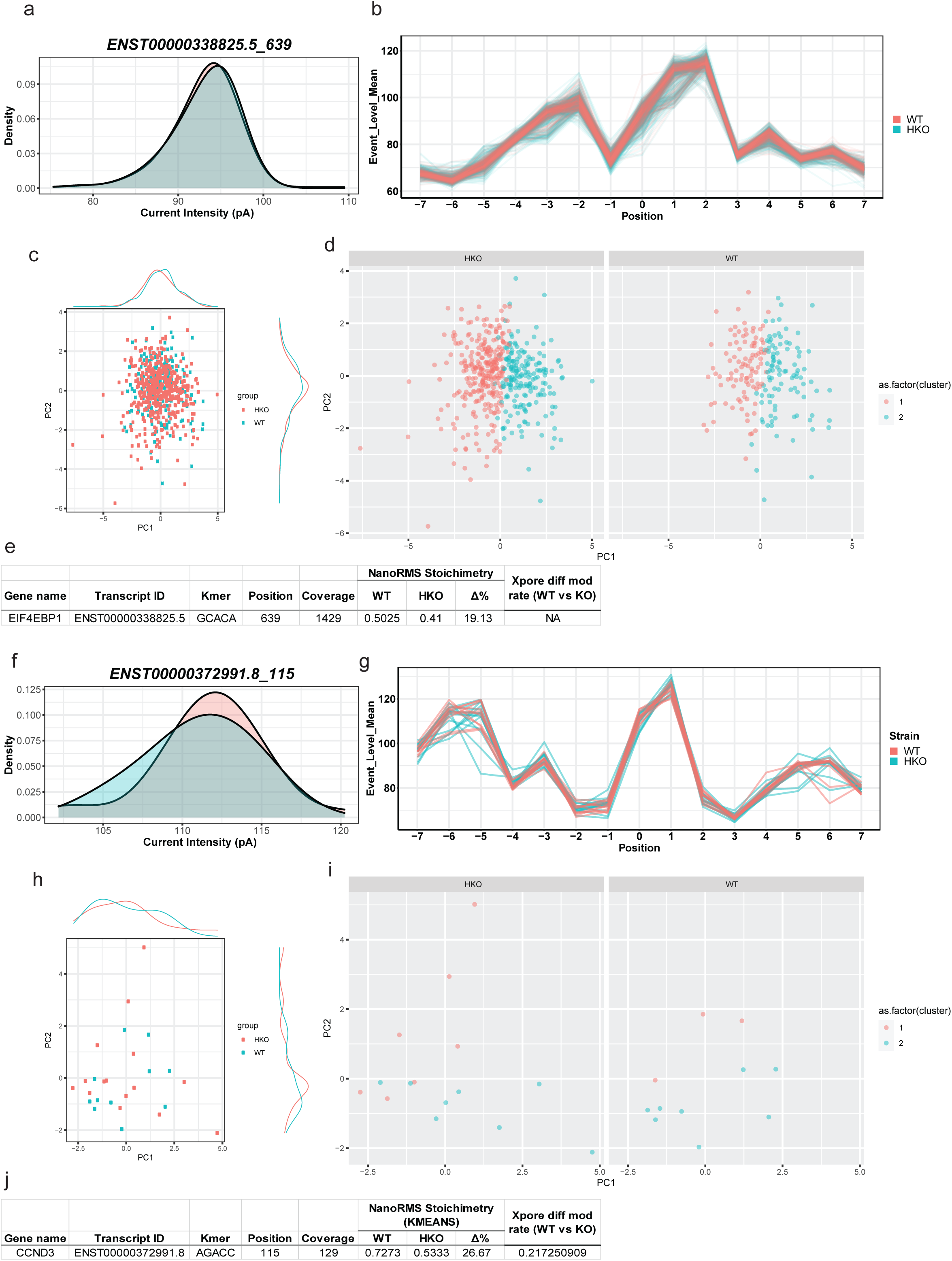
Heterozygous loss of METTL3 leads to differential methylation of DRACH motifs on EIF4EBP1 and Cyclin D3. Candidate m^6^A sites on EIF4EBP1 identified by nanocompore (KS intensity context 2) and CCND1 (Cyclin D3) mRNA identified by the XPore analysis tool was investigated using a modified nanoRMS pipeline. **a)** Current intensity Density plot of the 4EBP1 transcript ENST00000338825.5, showing the location of the differentially modified Adenosine at position 639. **b)** Per-read Current intensity analysis of WT and HKO reads in a 15bp window centred at position 639 of EIF4EBP1 (WT in red, HKO-1 in blue). **c)** PCA scatterplot for EIF4EBP1, position 639. Here only the first two principal components (PC1 and PC2) are shown. **d)** WT and HKO unsupervised KMEANS read clustering for EIF4EBP1 position 639. Factor 1 (red) is inferred to be m^6^A modified and factor 2 (blue) is inferred to be unmodified. **e)** WT and HKO KMEANS contingency table with NanoRMS calculated stoichiometry change of EIF4EBP1 position 639. **f)** Current density plot of of the CCND3 (Cyclin D3) transcript ENST00000372991.8 centred on the differentially modified Adenosine at position 115. **g)** Per-read Current intensity analysis of WT and HKO reads in a 15bp window centred at position 115 of CCND3 (WT in red, HKO-1 in blue). Principal component analysis (PCA) was carried out for each site using 15mer windows on current intensity values. **h)** PCA analysis carried out on position 115 of CCND3 using a 15mer on current intensity. Here only PCA1 and PCA2 are shown. **i)** WT and HKO unsupervised KMEANS read clustering for CCND3 at position 115. Factor 1 (red) is inferred to be m^6^A modified and factor 2 (blue) is inferred to be unmodified. **j)** WT and HKO KMEANS contingency table with NanoRMS calculated stoichiometry change of CCND3 position 115. XPore predicted differential modification rate is also included.

We next analysed a DRACH site in *EIF4EBP1* identified as differentially modified by Nanocompore (KS test on current intensity, context ±2, p < 0.01; Supplementary Tables 1 and 2). This transcript showed uniform coverage across its length, with a minimum depth exceeding 100 full-length reads (Supplementary Figure 6a,b). Analysis of current intensity revealed a modest shift at the DRACH site (position 639), as well as a pronounced shift at the immediately adjacent +1 position, suggesting that m6A may influence nanopore signal beyond the modified nucleotide itself (Figure 4a,b). Principal component analysis revealed partial separation of WT and HKO reads, indicating subtle but reproducible differences in signal features (Figure 4c). KMEANS clustering estimated an approximately 20% reduction in m6A stoichiometry at this site in HKO cells compared to WT (Figure 4d,e).

Together, these analyses demonstrate that nanoRMS-based stoichiometry estimates are consistent with differential modification rates inferred from xPore and Nanocompore and support the conclusion that partial loss of METTL3 leads to reproducible, site-specific reductions in m6A modification at transcripts involved in translation-related pathways.

### Partial loss of METTL3 is associated with increased translation and altered signalling

To determine how partial loss of METTL3 impacts protein regulation, we performed quantitative proteomic profiling of MDA-MB-468 cells following either genetic (METTL3 heterozygous knockout; HKO) or pharmacological (STM2457, 0.145 µM) reduction of METTL3. Both conditions resulted in widespread changes in protein abundance relative to WT cells (Fig. 5a,b), with substantial overlap between differentially expressed proteins (Fig. 5c). Hierarchical clustering revealed coordinated regulation of proteins across conditions (Fig. 5d), and pathway enrichment analysis identified processes related to cellular stress responses, metabolism, and growth-associated pathways (Fig. 5e).

**Figure 5.**
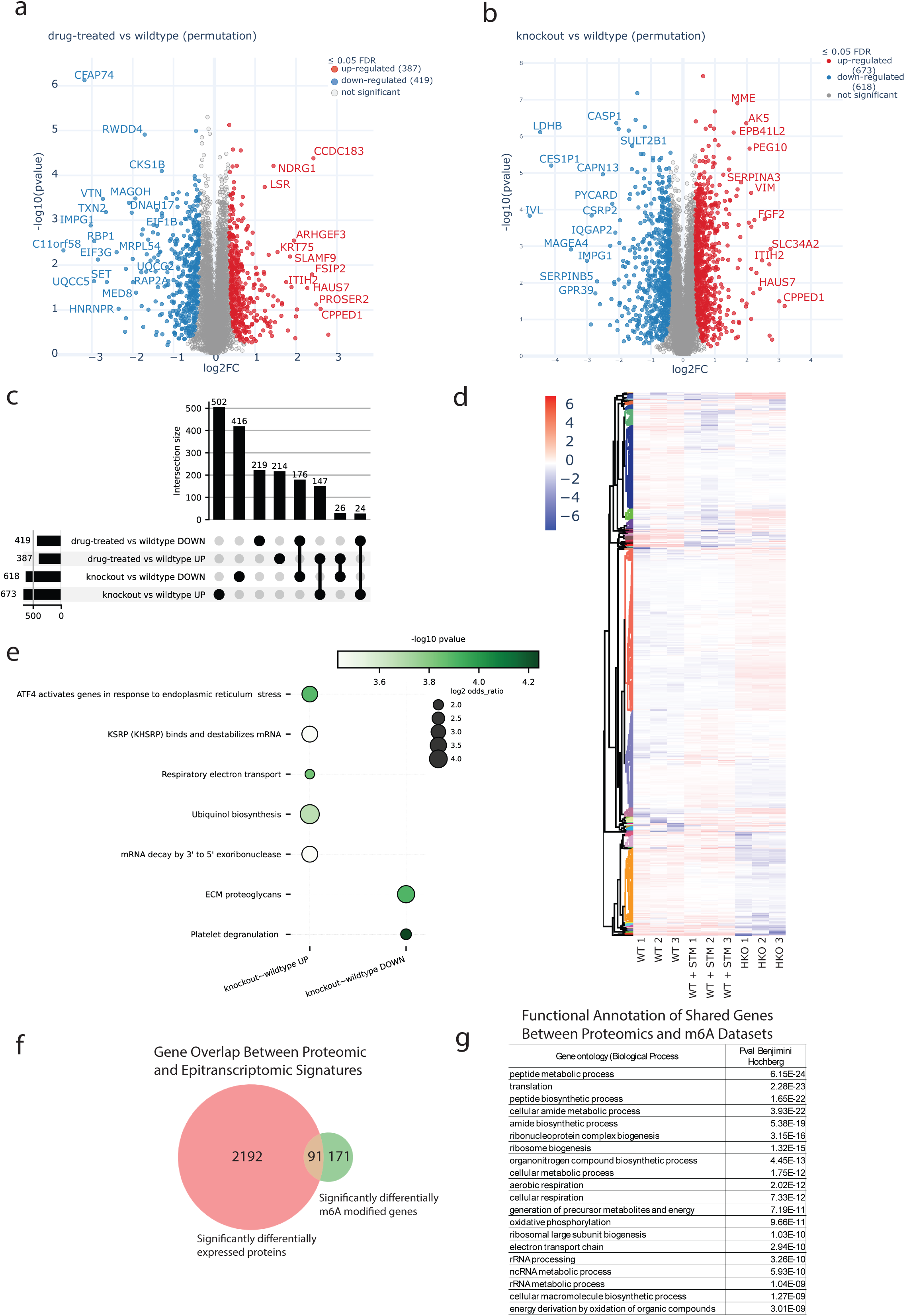
Partial reduction in METTL3 expression controls stress-response protein regulation independently of m6A deposition. **a)** Volcano plots of differential protein abundance in MDA-MB-468 TNBC cells treated with STM2457 (0.145::JµM, 72::Jh) vs WT (MinProb; permutation. FDR::J<::J0.05; *n*::J=::J3). **b)** Volcano plots for WT vs METTL3 heterozygous knockout (HKO) cells (as described above in **a)**. **c)** UpSet plot showing overlap of significantly differentially expressed proteins between drug-treated and METTL3 HKO cells. **d)** Heatmap of 2,097 significantly differentially expressed proteins (≥1 condition; FC::J>::J1; *p*::J<::J0.05); red::J=::Jincrease, blue::J=::Jdecrease. **e)** Reactome pathway enrichment of significantly regulated leading-razor proteins (adj. *p*::J<::J0.05; |log₂FC|::J≥::J1; Fisher’s exact test, perm. FDR::J<::J0.05). **f)** Venn diagram showing overlap of differentially methylated transcripts to steady-state protein changes (HKO vs WT). **g)** Top 20 enriched Gene Ontology (GO) biological process terms among genes showing both differential m6A methylation and protein expression changes.

Integration of proteomic data with m6A profiling demonstrated limited overlap between differentially methylated transcripts and changes in steady-state protein abundance (Fig. 5f,g), indicating that many protein-level changes observed following partial METTL3 reduction occur independently of direct alterations in m6A deposition.

To assess whether partial loss of METTL3 alters cellular signalling states, we next performed global phosphoproteomic profiling. Both STM2457-treated WT cells and METTL3 HKO cells exhibited extensive changes in protein phosphorylation relative to WT (Fig. 6a,b), with overlapping sets of differentially phosphorylated proteins across conditions (Fig. 6c). Pathway enrichment analysis of differentially phosphorylated proteins revealed increased representation of mTORC1-associated signalling pathways following partial reduction of METTL3 abundance or activity (Fig. 6e), consistent with altered regulation of growth and translational control pathways.

**Figure 6.**
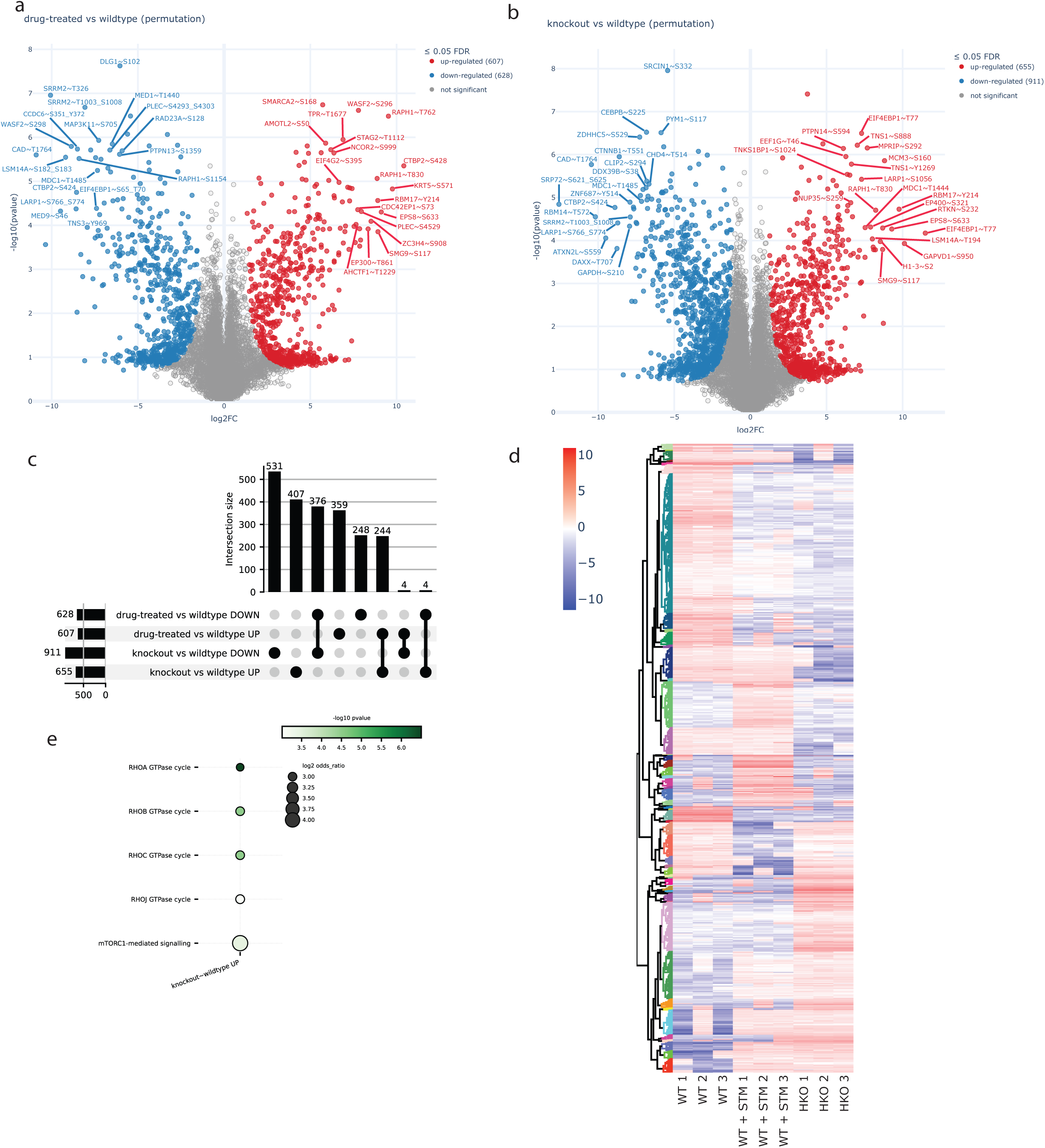
Reduction of m6A expression leads to increased mTORC1 signalling. **a)** Volcano plot showing differential phosphorylation of proteins in MDA-MB-468 cells treated with STM2457 (0.145::JµM, 72::Jh) compared to wild-type (WT) cells. Data were log₂-transformed, filtered for modified peptides, and imputed using MinProb (width = 0.3, shift = 1.8). Statistical analysis was performed using unpaired t-tests with permutation-based FDR correction (FDR < 0.05, fold change > 1, n = 3 biological replicates). **b)** Volcano plot comparing WT cells to METTL3 heterozygous knockout (HKO) cells under the same processing and statistical conditions as in **a)**. **c)** UpSet plot illustrating the overlap of significantly differentially phosphorylated proteins between STM2457-treated and METTL3 HKO cells. **d)** Heatmap of 2801 significantly differentially phosphorylated proteins detected by LC-MS in either STM2457-treated or METTL3 HKO cells compared to WT (significant in ≥1 condition; fold change > 1; p < 0.05). Red indicates increased phosphorylation; blue indicates decreased phosphorylation. All conditions include 3 biological replicates. **e)** Reactome pathway enrichment of proteins undergoing differential phosphorylation upon partial reduction of METTL3 abundance or activity. Only conditions with more than one significantly enriched ontology term (adjusted p-value < 0.05, ≥2 proteins per term) are shown.

To validate and further resolve these signalling changes, we examined key components of the PI3K/AKT/mTOR pathway by immunoblotting. Partial genetic or pharmacological reduction of METTL3 was associated with increased phosphorylation of mTOR at Ser2448 and elevated phosphorylation of AKT at Ser473, alongside increased abundance of PI3K (Fig. 7a). In contrast, ERK1/2 phosphorylation was largely unchanged, indicating pathway selectivity rather than global activation of mitogenic signalling.

**Figure 7.**
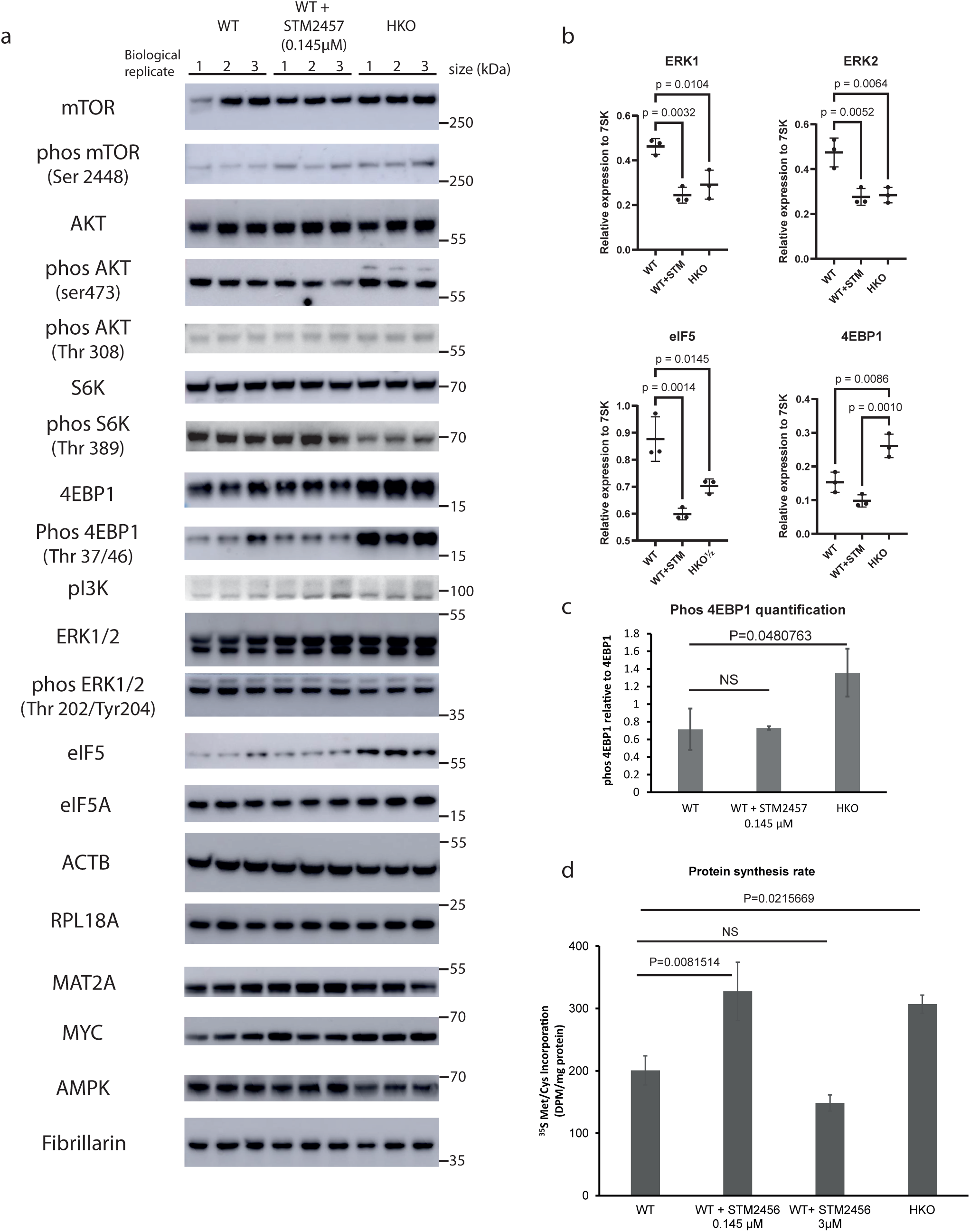
METTL3 regulates the PI3K/AKT/mTOR pathway. **a)** Western blot analysis of Akt-mTOR and ERK pathway activation in WT, 0.145 µM STM2457-treated WT, and HKO cell lines. Expression and phosphorylation changes in key components (Akt, mTOR, S6K, EIF4EBP, PI3K, ERK1/2) were assessed. Additionally, classical 5’TOP mRNA, RPL18A and non-5’ TOP translation factors (eIF5, eIF5A) were analysed^52,53^. ACTB and Fibrillarin served as loading controls. **b)** Quantification of ERK1, ERK2, eIF5, and 4EBP1 mRNA levels by qRT-PCR, comparing samples with or without METTL3 inhibition/knockout. Data represent the average (n=3) ± SEM, normalized to RN7SK (7SK). P-values were calculated using one-way ANOVA followed by Tukey HSD multiple comparison test. **c)** Relative phosphorylation of 4EBP1 (Thr 37/46) normalized to total 4EBP1 protein levels, as determined by Western blot analysis using Fiji^54^. **d)** Protein synthesis rates in STM2457 treated MDA-MB-468 cells and in the HKO knockout line as determined by [35S]-Met/Cys labelling. Disintegrations per minute (dpms) are graphed in the Y-axis.

Downstream of mTORC1, METTL3 HKO cells displayed increased steady-state levels and phosphorylation of EIF4EBP1 (Thr37/46), an effect that was also reflected at the transcript level (Fig. 7b,c). Despite this increase, phosphorylation of S6K was reduced in HKO cells, while remaining largely unchanged following low-dose STM2457 treatment (Fig. 7a). This divergence in downstream mTORC1 outputs suggests selective modulation of translational control mechanisms rather than uniform pathway activation.

Consistent with these signalling changes, partial genetic or pharmacological reduction of METTL3 resulted in increased global protein synthesis, as measured by [³::JS]-methionine/cysteine incorporation (Fig. 7d). In contrast, stronger METTL3 inhibition suppressed protein synthesis, paralleling the reduced proliferative capacity observed under these conditions (Fig. 2).

Together, these data demonstrate that partial loss of METTL3 leads to increased translational output accompanied by selective rewiring of growth-associated signalling pathways. These effects are not fully explained by changes in m6A deposition alone, highlighting a dosage-sensitive role for METTL3 in coordinating translational capacity and signalling state.

## Discussion

The consequences of m6A modification on mRNA are varied and complex, affecting all aspects of RNA metabolism, including but not limited to RNA stability, splicing, localisation, and translation. As such, changes in m6A deposition can have a wide range of effects on cellular processes and must be tightly regulated. In this study, we show that heterozygous loss of METTL3, and thus a modest reduction in m6A deposition on mRNA, leads to activation of the PI3K/AKT/mTORC1 signalling pathway, resulting in increased translation and proliferation. This is consistent with previously published studies in TNBC and several other cancers, where partial loss of METTL3 expression is associated with poor prognosis. We find that a relatively small reduction in m6A preferentially affects transcripts involved in translation and metabolism, whereas strong reduction of METTL3 activity or expression inhibits both proliferation and translation.

Insulin treatment increases proliferation in cells treated with 3 µM STM2120 and 0.145 µM STM2457, but not in METTL3 heterozygous knockout cells or cells treated with 3 µM STM2457. Rapamycin treatment results in a pronounced inhibition of growth across all genotypes and METTL3 inhibitor treatments, whereas metformin, which activates AMPK and thereby inhibits mTORC1, does not significantly affect proliferation, excluding AMPK as a major contributor to the observed phenotype.

EGF treatment results in the most profound inhibition of cell growth in WT cells and all METTL3 inhibitor–treated lines, with the exception of the METTL3 heterozygous knockout cell line. This is consistent with EGF-mediated senescence and apoptosis previously reported in MDA-MB-468 cells due to their exceptionally high levels of EGFR expression. Dexamethasone treatment, which has been shown to counteract EGF-mediated senescence and apoptosis, does not rescue proliferation, except in the METTL3 heterozygous knockout cell line. EGFR overexpression is a hallmark of TNBC and numerous other cancers and is associated with poor prognosis. In MDA-MB-468 cells, this extremely high EGFR abundance induces apoptosis upon EGF stimulation, as previously described. ^46^

Differential methylation analysis using direct RNA sequencing reveals that partial loss of METTL3 leads to depletion of a subset of m6A sites, with reductions occurring across all DRACH motifs. This suggests that reduced METTL3 activity leads to a relatively uniform decrease in m6A deposition, regardless of the intrinsic modification frequency of individual DRACH sequences. This has implications for understanding the mechanisms governing differential m6A deposition across the transcriptome.

The distribution of differentially modified sites identified by both Nanocompore and xPore is broadly similar, with the highest proportion of sites located just upstream of the stop codon. Differences between the methods include a higher density of differentially methylated sites across the coding sequence detected by xPore and greater enrichment toward the end of the 3′UTR detected by Nanocompore. Notably, this distribution does not match predictions from m6Anet or patterns reported in previous studies, where m6A sites are most enriched within the 3′UTR, with the highest peak located on or immediately downstream of the stop codon. ^3,47,48^

These findings suggest that the sites most sensitive to METTL3 loss are located outside of canonical 3′UTR hotspots, particularly in regions immediately upstream of the stop codon.

Taken together, these results indicate that transcriptomic position, rather than DRACH motif strength, plays a dominant role in determining m6A deposition frequency. This conclusion is consistent with studies demonstrating that m6A is enriched at positions located more than 100 nucleotides from splice junctions and proximal to stop codons.^48–50^

Accumulating evidence suggests that m6A RNA modification and mTOR signalling are intrinsically linked.^45^ In our study, heterozygous knockout of METTL3, or treatment with low concentrations of the METTL3 inhibitor STM2457, result in increased expression of PI3K and 4EBP1, as well as increased phosphorylation of Akt (Ser473), mTOR (Ser2448), and 4EBP1 (Thr37/46). We further observe that heterozygous loss of METTL3 increases total 4EBP1 expression and phosphorylation, without a proportional increase in phosphorylation ratio, and leads to reduced phosphorylation of S6K. Reduced S6K phosphorylation upon METTL3 depletion has also been reported in the breast cancer cell lines MDA-MB-231 and T47D following METTL3 knockdown. ^51^

Nanocompore analysis identified a single significantly differentially modified DRACH motif within the 4EBP1 transcript, which may contribute to the observed increase in RNA and protein abundance following METTL3 depletion. However, this modification was not detected using the xPore pipeline, and pharmacologic inhibition of METTL3 did not reproduce the increase in 4EBP1 expression, suggesting that chronic METTL3 reduction or additional downstream targets may be required to drive this response.

Overall, our findings demonstrate that cellular growth and translational efficiency are highly sensitive to modest changes in m6A levels. Partial loss of METTL3 enhances proliferation and protein synthesis, consistent with deregulated METTL3 expression observed in TNBC, whereas strong inhibition of METTL3 produces a marked suppression of both processes. These data underscore the importance of maintaining m6A homeostasis for normal cellular function and raise important considerations regarding the therapeutic targeting of METTL3.

Notably, transcripts encoding proteins involved in translation are disproportionately affected by partial loss of m6A, implying a direct and selective impact on growth-related pathways. Our data suggest that components of the mTOR signalling axis are particularly sensitive to reduced METTL3 activity, providing a mechanistic link between METTL3 deregulation and enhanced proliferative capacity in cancer. While current therapeutic strategies largely focus on complete inhibition of METTL3, our findings suggest that restoring or stabilising METTL3 expression in contexts of partial loss may represent an alternative strategy to re-establish translational control and limit tumour growth.

## Methods

### Cell culture

Unless stated otherwise, MDA-MB-468 (obtained from ATCC (HTB-132)) and heterozygous knockout cell lines were maintained in DMEM containing 4.5g/L-glucose, L-Glutamine and sodium pyruvate (Thermo Fisher 41966-029), supplemented with 10% (v/v) foetal bovine serum and 1 % (v/v) penicillin/streptomycin at 37°C.

### Transfection and genotyping of knockout lines

Plasmids were transfected in the MDA-MB-468 cell line using FuGENE HD (Promega) non-liposomic transfection reagent. 5’ and 3’ sgRNA encoding plasmids (2 µg each) were co-transfected in 1:3 ratio with FuGENE reagent. Transfection medium was replaced after 24 h incubation. 1-2 days after cells were recovered from transfection, puromycin selection (0.7 µg/ml) was started for 24 hours. Selected cells were grown in low density for colony formation. Colonies were then extracted and isolated. After sufficient colony size, genomic DNA was extracted with GeneJET Genomic DNA Purification Kit (K0722, Thermo Fischer Scientific), and subjected to PCR [1X PCR buffer; 1.5 mM MgCl2; 0.2 mM dNTP; 0.5 µM primers; 50 ng genomic DNA; 1 U Taq DNA polymerase (10342020, Thermo Fischer Scientific)]. PCR products were separated on 1% agarose-TBE gel. Genomic DNA bands were extracted from agarose for Sanger sequencing.

### Generation of METTL3 heterozygous knockout cells

Heterozygous *METTL3* knockout in MDA-MB-468 cells was achieved using the CRISPR-Cas9 system, according to the published protocol by the Zhang lab (Ran et al., 2013). pSpCas9(BB)-2A-Puro (PX459) V2.0 was a gift from Feng Zhang (Addgene plasmid 62988; http://n2t.net/addgene:62988; RRID:Addgene 62988). The plasmid encodes the sgRNA sequence, Cas9 nuclease, and puromycin resistance gene for selection. sgRNA sequences were designed to disrupt the first exon of *METTL3*. The 5’ guide RNA (5’ CTTCCAAGAAAGCGCGACAC 3’) was designed to target 155 bp upstream of the translation start codon ATG. The 3’ sgRNA (5’ AGAGGGAGTACCAATGTACG 3’) targeted the Cas9 nuclease 149 bp downstream of the first exon. Control primers for genomic DNA PCR were designed to surround the deletion site (FW 5’ CCAACACCTAACGTGGCTCT 3’; RV 5’ ACAGCTGCCAAGAAATGACC 3’), amplifying 712 bp region. The sgRNA sequences were designed as complementary oligonucleotides with BbsI restriction enzyme sites attached. This allowed for one-step cloning and ligation of the annealed sgRNAs into the PX459 vector.

### RNA extraction RT/qRT-PCR analysis

Total RNA was extracted using Trizol (Invitrogen 15596026) according to the manufacturers protocol and reverse transcribed using RevertAid reverse transcription kit (Thermo Fisher K16191) using random primers and analysed by qPCR in an AriaMx PCR system (Agilent) using the Platinum SYBR Green qPCR SuperMix-UDG kit (Thermo Fisher 11733046). Oligonucleotide sequences used are detailed in supplementary table S5) qPCR analysis was carried out using the 2-ΔCt method by normalising against RN7SK or GAPDH levels.

### Polyadenylated transcript selection

Purified RNA for RNA mass spectrometry and Nanopore Direct RNA sequencing was subjected to poly(A) purification using either the Lexogen Poly(A) RNA selection kit V1.5 (157.96) or the NEBNext Poly(A) mRNA Magnetic Isolation Module (E7490L) according to the manufacturer’s instructions. To assess poly(A) selection efficiency, we employed the Fragment Analyzer system (Agilent) with the HS RNA kit (15nt) (Agilent DNF-472) using the mRNA preset program.

### RNA Mass spec

The RNA extraction was carried out using Trizol as detailed above, following the manufacturer’s protocol. However, instead of sodium acetate, ammonium acetate was used for precipitation. RNA was hydrolyzed to ribonucleosides by 20 U benzonase (Santa Cruz Biotech) and 0.2 U nuclease P1 (Sigma-Aldrich, Saint-Louis, MO) in 10 mM ammonium acetate pH 6.0 and 1 mM magnesium chloride at 40 °C for 1 hour, then added ammonium bicarbonate to 50 mM, 0.002 U phosphodiesterase I and 0.1 U alkaline phosphatase (Sigma-Aldrich, Saint-Louis, MO) and incubated further at 37 °C for 1 hour. The hydrolysates were added 3 volumes of acetonitrile and centrifuged (16,000 g, 30 min, 4 °C). The supernatants were lyophilized and dissolved in 50 µl water for LC-MS/MS analysis of modified and unmodified ribonucleosides. Chromatographic separation was performed using an Agilent 1290 Infinity II UHPLC system with an ZORBAX RRHD Eclipse Plus C18 150 x 2.1 mm ID (1.8 μm) column protected with an ZORBAX RRHD Eclipse Plus C18 5 x 2.1 mm ID (1.8 µm) guard column (Agilent Technologies, Foster City, CA). The mobile phase consisted of water and methanol (both added 0.1% formic acid) run at 0.23 ml/min, for modifications starting with 5% methanol for 0.5 min followed by a 2.5 min gradient of 5-15 % methanol, a 3 min gradient of 15-95% methanol and 4 min re-equilibration with 5% methanol. A portion of each sample was diluted for the analysis of unmodified ribonucleosides which was chromatographed isocratically with 20% methanol. Mass spectrometric detection was performed using an Agilent 6495 Triple Quadrupole system with electrospray ionization, monitoring the mass transitions 268.1-136.1 (A), 284.1-152.1 (G), 244.1-112.1 (C), 245.1-113.1 (U), and 282.1-150.1 (m6A) in positive ionization mode.

### Proteomics data processing and statistical analysis

Global proteome and phosphoproteome datasets were processed by the Proteomics Research Infrastructure (PRI, University of Copenhagen) using the Proteomelit analysis pipeline (v1.1.0), adapted from the Clinical Knowledge Graph workflow. Protein and modified peptide intensities were log2-transformed and filtered to retain features with at least two valid values in at least one experimental group. Missing values were imputed using either MinProb (width = 0.3, shift = 1.8) or a mixed imputation strategy in which missing-at-random values were imputed using k-nearest neighbours and missing-not-at-random values using MinProb. For phosphoproteomics, precursor intensities were aggregated to modified peptides, filtered to retain phosphopeptides, and normalised to the corresponding protein abundance measured in the global proteome dataset.

Differential abundance between conditions (drug-treated vs wildtype and knockout vs wildtype) was assessed using unpaired t-tests with multiple hypothesis correction using the Benjamini–Hochberg procedure and permutation-based false discovery rate (FDR) estimation (250 permutations). Functional enrichment analyses were performed using Fisher’s exact test with Benjamini–Hochberg correction using Gene Ontology and Reactome annotations. Kinase–substrate and motif enrichment analyses for the phosphoproteome were performed using annotations from PhosphoSitePlus and Phosida.

### Nanopore direct RNA sequencing

500 ng of purified, polyadenylated RNA from WT and HKO were prepared as described above. Direct RNA sequencing libraries were prepared using the Nanopore Direct RNA sequencing kit (SQK-RNA002) as directed. Sequencing was performed using R9.4.1 sequencing chemistry in FLO-MIN106 flow cells for a period of 72h or until pore exhaustion, which typically took between 12-48h.

### Cell lysis and western blotting

Cells were treated with or without the indicated concentrations of STM2457 (Storm Therapeutics, Medchem Express, HY-134836) for 24 hours. They were then pelleted and lysed in RIPA buffer (50 mM Tris–HCl, pH 7.5, 1% Triton X-100, 0.5% Na-deoxycholate, 0.1% SDS, 150 mM NaCl, 1mM EDTA), supplemented with 1mM PMSF, cOmplete EDTA-free protease inhibitor cocktail (Roche 04693159001), and PhosSTOP phosphatase inhibitor (Roche PHOSS-RO). The lysate was cleared by centrifugation.

Protein concentrations were determined by Bicinchoninic acid assay (BCA) using the Pierce BCA Protein Assay Reducing Agent Compatible kit (Pierce, Thermo Fisher 23250) and quantified on a Glomax Discover microplate reader (Promega).

Lysates were mixed with NuPage LDS sample buffer (Novex NP0007) and reducing agent (Novex NP0009), then denatured at 80°C for 10 minutes. Depending on the target protein size, the samples were loaded onto either Novex Nupage 4-12% Bis-Tris gels (Invitrogen) or Novex Nupage 3-8% Tris acetate gels (Invitrogen). The Bis-Tris gels were run with either Nupage MOPS SDS (Novex NP0001) or Bolt MES SDS (Novex B0002) running buffer. The Tris acetate gels were run with Tris tricine SDS running buffer (10mM Tris, 10mM Tricine, 0.01% SDS, pH8.3). Gels were transferred onto a PVDF membrane using either the iBlot2 system (Thermo Fisher) or a wet transfer onto Immobilon-P PVDF membrane (0.45µm pore size, Millipore IPVH0005) and Novex transfer buffer (NP0006-1), supplemented with 20% methanol.

Membranes were blocked with PBS-0.05% Tween and 5% milk for 1 hour at room temperature with agitation, before overnight incubation with primary antibodies. Proteins were visualized using SuperSignal West Pico PLUS ECL (Pierce 34580) or SuperSignal West Femto Maximum Sensitivity Substrate (Pierce 34094) on a GE Amersham Imager 600.

Antibodies used in this study and the dilutions used are detailed in supplementary table 2. Protein bands were quantified using Image J and normalised to either Fibrillarin or Beta-Actin.

### Proliferation capacity assays

MDA-MB-468 triple-negative breast cancer cells were cultured at 37 °C and 5% CO₂ and seeded at 3,000 cells per well in 96-well plates. After overnight attachment, cells were pre-treated for 24 h with STM2457 (0.145 µM or 3 µM) or the weak METTL3 inhibitor analogue STM2120 (3 µM).

Cells were then exposed to EGF, EGF plus dexamethasone, insulin, rapamycin, metformin, or left untreated under glucose concentrations of 0, 5.5, 15, or 25 mM, with media refreshed at 72 h. Proliferation capacity was assessed by CellTiter-Glo luminescence at 24, 48, 72, 96, and 120 h using separate plates for each time point, with background subtraction.

Luminescence data were summarized primarily as absolute area under the curve (AUC; 24–120 h) using trapezoidal integration. Secondary analyses included normalized trajectories (each series divided by its 24 h value; time rebased so 24 h = 0 h), normalized AUC (0–96 h), windowed signal change rates (log₂/h), maximum signal, and time to half-maximal signal (T₅₀). Three biological replicates were analysed; replicate-level endpoints were compared to WT using Welch’s t-tests with Benjamini–Hochberg FDR correction, and effect sizes were reported as Cohen’s d. Data are shown as vertical heatmaps or mean ± SEM luminescence trajectories, with normalized dynamics presented in the Supplementary Information.

### Measurement of de novo protein synthesis by [^35^S]-Met/Cys labelling

WT and HKO cells were seeded at a concentration of 7 x10^5^ cells per well. After 24h WT cells were treated either with or without the indicated concentrations of STM2457. After a further 24h, the cells were washed and starved in Cysteine and Methionine-free DMEM (21013024) supplemented with 10% FBS, 1% Penicillin/Streptomycin and 2mM L-Glutamine for 3 hours. They were then pulsed with 30 µCi/mL of EasyTag EXPRESS^35^S Protein Labelling Mix, [^35^S] (Perkin Elmer/Revity NEG772014MC) for 4 hours. In order to arrest translation, cells were treated with 50ng/mL of cycloheximide (Merck C4859-1ML) for 5 min. Cells were then immediately lysed in ice cold RIPA buffer and quantified using BCA as detailed above. ^35^S Met and Cys incorporation was then measured using a Perkin-Elmer Tri-Carb 4910 TR scintillation counter.

### Bioinformatics analysis

#### Nanopore Direct RNA Sequencing Data processing

Multi-fast5 files were basecalled using Guppy v4.0.11 using the rna_r9.4.1_70bps_hac.cfg high accuracy RNA basecalling model and were basecalled on the GenomeDK HPC using two Nvidia V100 16Gb GPUs. In addition, the --fast5_out option was used to allow storing of trace tables. Fastq files were indexed with nanopolish and reads aligned to the Gencode v33 transcriptome using the -x map-ont preset. Mapped reads were then filtered using samtools view using the -F 2324 flagstat and a minimum map quality of 10. Following this the reads were polished using nanopolish eventalign using the --scale-events and --signal-index, and--print-read-names options.

#### Nanocompore v1.04

Following read resquiggling by nanopolish, the output was passed to NanopolishComp Eventalign_collapse to collapse the data per kmer rather than by event. The nanocompore SampComp module was then run using the following parameters, logit =TRUE, min_coverage =30, bed_fn, comparison_methods= [“GMM”, “KS”, “MW”, “TT”], sequence_context=2, sequence_context_weights= “harmonic”. For nanocompore WT replicate 1, 2 and 3 was compared to HKO replicate 2 and 3. This was done as these replicates has a more similar depth. In order to improve performance for very highly abundant transcripts, SampComp was run with downsample-high-coverage = 5000.

#### XPore v2.0

Using the output of nanopolish eventalign above we ran the dataprep module to prepare the data for analysis. Following the datapre step, the Diffmod module was run with the following configuration criteria, readcount_min=15, readcount_max=1000, prefiltering method= t-test, prefiltering threshold=0.1, max_iters=500 and stopping criteria=0.00001. Finally, to consider single modification types per kmer, we ran the spore postprocessing module.

#### M6Anet v2.0.2

After the eventalign step described in the DRS data processing section above, the data was first preprocessed using the m6anet dataprep module after which we ran m6anet inference with the number of iterations set to 1000. With this m6A net samples 20 reads per site and average the probability that a site is modified 1000 times. As m6Anet predicts RNA modifications using a multiple instance learning model and does not support differential modification analysis, m6A net was run for each cell line individually, pooling over multiple replicates.

### NanoRMS

#### Identification of candidate modification sites

nanoRMS supports de novo and paired sample RNA modification prediction of Pseudouridine, through its characteristic C to U mismatch pattern. Such a pattern was not identified for m6A, and as such site identification was performed on the outputs of xPore. Statistically significant sites were identified using the filter pval <0.05 and z score >1

#### Extraction of Current-features

nanoRMS employs two separate pipelines for extracting current-features, Nanopolish and Tombo. Nanopolish allows extraction of Current intensities per read and position. Tombo however, allows for extraction of current intensity as well as dwell times and the nanoRMS script get_features allows for integration with trace data.

We utilised a modified python translation of the nanoRMS script nanopolish_window.R (windowscript.py) that creates 15-mer windows centred on previously identified candidate positions with the collapsed reads. This script can handle much larger datasets than the original R script, with minimal memory use.

Finally we modified the nanoRMS script nanopoilish_export_each_position.R (nanopolish_export_each_position_singleinput.R). to sort these windows by individual reads and export them for clustering. The modification consists of producing windows for specific positions instead of all positions in a given dataset.

Estimation of stoichiometry was done by KMEANS clustering with the script read_clustering_fixed.R. which implements a modification in the input handling, of the the original nanoRMS script, read_clustering.R, to produce a contingency table with the summarized clusters and graphs of these clusters.

#### Functional analysis of differentially modified genes

Functional analysis of significantly differentially modified genes based on the GO, KEGG and REACTOME databases (Gene Ontology; http://www.geneontology.org, Kyoto Encyclopaedia of genes and Genomes, http://www.genome.jp/kegg/ and http://reactome.org) was implemented using the humanmine webtool (Kalderimis et al. 2014). GO, KEGG and REACTOME enriched terms were selected using a False discovery rate of <0.05 (Holm-Bonferroni correction).

#### Metagene plots

Metagene plots were created using the Guitar R package and the TxDb.Hsapiens.UCSC.hg.38.knownGene package. The TxDb package provided the genomic TxDb object required to assign mRNA landmarks. Plots were made with the GuitarPlot function using the pltTxtype = c(“mrna “) aurgument.

## Data availability

The RNA sequencing data can be accessed at GEO GSE272282.

The code can be accessed at https://doi.org/10.5281/zenodo.12530077 and https://github.com/samir-watson/METTL3-HKO-paper-code

## Acknowledgments

Work in the author’s lab is funded by the Lundbeck Foundation, the Novo Nordisk Foundation, Danish Cancer Association, the Carlsberg Foundation, Independent Research Fund Denmark, Aage and Johanne Louis-Hansen’s Foundation.

## Author contributions

SW designed experiments, performed experiments and data analysis, interpreted data, wrote the manuscript, approved final manuscript.

JL designed experiments, performed experiments, interpreted data, approved final manuscript. SNK performed experiments, interpreted data, approved final manuscript.

PAD performed data analysis and approved the final manuscript.

AB performed RNA mass spec experiments and interpreted data. Approved the final manuscript.

HOP performed experiments and approved the final manuscript.

UAVØ designed experiments, interpreted data, supervised work, secured funding, wrote the manuscript, approved final manuscript.

## References

1. Wei, C.-M., Gershowitz, A., and Moss, B. (1975). Methylated nucleotides block 5′ terminus of HeLa cell messenger RNA. Cell 4, 379–386. 10.1016/0092-8674(75)90158-0.

2. Workman, R.E., Tang, A.D., Tang, P.S., Jain, M., Tyson, J.R., Razaghi, R., Zuzarte, P.C., Gilpatrick, T., Payne, A., Quick, J., et al. (2019). Nanopore native RNA sequencing of a human poly(A) transcriptome. Nat. Methods 16, 1297–1305. 10.1038/s41592-019-0617-2.

3. Dominissini, D., Moshitch-Moshkovitz, S., Schwartz, S., Salmon-Divon, M., Ungar, L., Osenberg, S., Cesarkas, K., Jacob-Hirsch, J., Amariglio, N., Kupiec, M., et al. (2012). Topology of the human and mouse m6A RNA methylomes revealed by m6A-seq. Nature 485, 201–206. 10.1038/nature11112.

4. He, P.C., and He, C. (2021). m6A RNA methylation: from mechanisms to therapeutic potential. EMBO J. 40, e105977. 10.15252/embj.2020105977.

5. Jiang, X., Liu, B., Nie, Z., Duan, L., Xiong, Q., Jin, Z., Yang, C., and Chen, Y. (2021). The role of m6A modification in the biological functions and diseases. Signal Transduct. Target. Ther. 6, 74. 10.1038/s41392-020-00450-x.

6. Meyer, K.D., Saletore, Y., Zumbo, P., Elemento, O., Mason, C.E., and Jaffrey, S.R. (2012). Comprehensive Analysis of mRNA Methylation Reveals Enrichment in 3′ UTRs and near Stop Codons. Cell 149, 1635–1646. 10.1016/j.cell.2012.05.003.

7. Jia, G., Fu, Y., Zhao, X., Dai, Q., Zheng, G., Yang, Y., Yi, C., Lindahl, T., Pan, T., Yang, Y.-G., et al. (2011). N6-Methyladenosine in nuclear RNA is a major substrate of the obesity-associated FTO. Nat. Chem. Biol. 7, 885–887. 10.1038/nchembio.687.

8. Zheng, G., Dahl, J.A., Niu, Y., Fedorcsak, P., Huang, C.-M., Li, C.J., Vågbø, C.B., Shi, Y., Wang, W.-L., Song, S.-H., et al. (2013). ALKBH5 Is a Mammalian RNA Demethylase that Impacts RNA Metabolism and Mouse Fertility. Mol. Cell 49, 18–29. 10.1016/j.molcel.2012.10.015.

9. Huang, H., Weng, H., Sun, W., Qin, X., Shi, H., Wu, H., Zhao, B.S., Mesquita, A., Liu, C., Yuan, C.L., et al. (2018). Recognition of RNA N6-methyladenosine by IGF2BP proteins enhances mRNA stability and translation. Nat. Cell Biol. 20, 285–295. 10.1038/s41556-018-0045-z.

10. Xiao, W., Adhikari, S., Dahal, U., Chen, Y.-S., Hao, Y.-J., Sun, B.-F., Sun, H.-Y., Li, A., Ping, X.-L., Lai, W.-Y., et al. (2016). Nuclear m6A Reader YTHDC1 Regulates mRNA Splicing. Mol. Cell 61, 507–519. 10.1016/j.molcel.2016.01.012.

11. Spitale, R.C., Flynn, R.A., Zhang, Q.C., Crisalli, P., Lee, B., Jung, J.-W., Kuchelmeister, H.Y., Batista, P.J., Torre, E.A., Kool, E.T., et al. (2015). Structural imprints in vivo decode RNA regulatory mechanisms. Nature 519, 486–490. 10.1038/nature14263.

12. Roundtree, I.A., Luo, G.-Z., Zhang, Z., Wang, X., Zhou, T., Cui, Y., Sha, J., Huang, X., Guerrero, L., Xie, P., et al. (2017). YTHDC1 mediates nuclear export of N6-methyladenosine methylated mRNAs. eLife 6, e31311. 10.7554/elife.31311.

13. Wang, X., Zhao, B.S., Roundtree, I.A., Lu, Z., Han, D., Ma, H., Weng, X., Chen, K., Shi, H., and He, C. (2015). N6-methyladenosine modulates messenger RNA translation efficiency. Cell. 10.1016/j.cell.2015.05.014.

14. Alarcón, C.R., Lee, H., Goodarzi, H., Halberg, N., and Tavazoie, S.F. (2015). N6-methyladenosine marks primary microRNAs for processing. Nature 519, 482–485. 10.1038/nature14281.

15. Wu, B., Su, S., Patil, D.P., Liu, H., Gan, J., Jaffrey, S.R., and Ma, J. (2018). Molecular basis for the specific and multivariant recognitions of RNA substrates by human hnRNP A2/B1. Nat. Commun. 9, 420. 10.1038/s41467-017-02770-z.

16. Geula, S., Moshitch-Moshkovitz, S., Dominissini, D., Mansour, A.A., Kol, N., Salmon-Divon, M., Hershkovitz, V., Peer, E., Mor, N., Manor, Y.S., et al. (2015). m6A mRNA methylation facilitates resolution of naïve pluripotency toward differentiation. Science 347, 1002–1006. 10.1126/science.1261417.

17. Pan, Y., Chen, H., Zhang, X., Liu, W., Ding, Y., Huang, D., Zhai, J., Wei, W., Wen, J., Chen, D., et al. (2023). METTL3 drives NAFLD-related hepatocellular carcinoma and is a therapeutic target for boosting immunotherapy. Cell Rep. Med. 4, 101144. 10.1016/j.xcrm.2023.101144.

18. Barbieri, I., Tzelepis, K., Pandolfini, L., Shi, J., Millán-Zambrano, G., Robson, S.C., Aspris, D., Migliori, V., Bannister, A.J., Han, N., et al. (2017). Promoter-bound METTL3 maintains myeloid leukaemia by m6A-dependent translation control. Nature 552, 126–131. 10.1038/nature24678.

19. Poh, H.X., Mirza, A.H., Pickering, B.F., and Jaffrey, S.R. (2022). Alternative splicing of METTL3 explains apparently METTL3-independent m6A modifications in mRNA. PLoS Biol. 20, e3001683. 10.1371/journal.pbio.3001683.

20. Batista, P.J., Molinie, B., Wang, J., Qu, K., Zhang, J., Li, L., Bouley, D.M., Lujan, E., Haddad, B., Daneshvar, K., et al. (2014). m6A RNA Modification Controls Cell Fate Transition in Mammalian Embryonic Stem Cells. Cell Stem Cell 15, 707–719. 10.1016/j.stem.2014.09.019.

21. Kim, K.L., van Galen, P., Hovestadt, V., Rahme, G.J., Andreishcheva, E.N., Shinde, A., Gaskell, E., Jones, D.R., Shema, E., and Bernstein, B.E. (2021). Systematic detection of m6A-modified transcripts at single-molecule and single-cell resolution. Cell Rep. Methods 1, 100061. 10.1016/j.crmeth.2021.100061.

22. Li, H.-B., Tong, J., Zhu, S., Batista, P.J., Duffy, E.E., Zhao, J., Bailis, W., Cao, G., Kroehling, L., Chen, Y., et al. (2017). m6A mRNA methylation controls T cell homeostasis by targeting the IL-7/STAT5/SOCS pathways. Nature 548, 338–342. 10.1038/nature23450.

23. Pomaville, M., Chennakesavalu, M., Wang, P., Jiang, Z., Sun, H.-L., Ren, P., Borchert, R., Gupta, V., Ye, C., Ge, R., et al. (2024). Small-molecule inhibition of the METTL3/METTL14 complex suppresses neuroblastoma tumor growth and promotes differentiation. Cell Rep. 43, 114165. 10.1016/j.celrep.2024.114165.

24. Yuan, Y., Du, Y., Wang, L., and Liu, X. (2020). The M6A methyltransferase METTL3 promotes the development and progression of prostate carcinoma via mediating MYC methylation. J. Cancer 11, 3588–3595. 10.7150/jca.42338.

25. Li, J., Rao, B., Yang, J., Liu, L., Huang, M., Liu, X., Cui, G., Li, C., Han, Q., Yang, H., et al. (2020). Dysregulated m6A-Related Regulators Are Associated With Tumor Metastasis and Poor Prognosis in Osteosarcoma. Front. Oncol. 10, 769. 10.3389/fonc.2020.00769.

26. Deng, X., Qing, Y., Horne, D., Huang, H., and Chen, J. (2023). The roles and implications of RNA m6A modification in cancer. Nat. Rev. Clin. Oncol. 20, 507–526. 10.1038/s41571-023-00774-x.

27. Shi, Y., Zheng, C., Jin, Y., Bao, B., Wang, D., Hou, K., Feng, J., Tang, S., Qu, X., Liu, Y., et al. (2020). Reduced Expression of METTL3 Promotes Metastasis of Triple-Negative Breast Cancer by m6A Methylation-Mediated COL3A1 Up-Regulation. Front. Oncol. 10, 1126. 10.3389/fonc.2020.01126.

28. Perou, C.M., Sørlie, T., Eisen, M.B., van de Rijn, M., Jeffrey, S.S., Rees, C.A., Pollack, J.R., Ross, D.T., Johnsen, H., Akslen, L.A., et al. (2000). Molecular portraits of human breast tumours. Nature 406, 747–752. 10.1038/35021093.

29. Nedeljković, M., and Damjanović, A. (2019). Mechanisms of Chemotherapy Resistance in Triple-Negative Breast Cancer—How We Can Rise to the Challenge. Cells 8, 957. 10.3390/cells8090957.

30. Garrido-Castro, A.C., Lin, N.U., and Polyak, K. (2019). Insights into Molecular Classifications of Triple-Negative Breast Cancer: Improving Patient Selection for Treatment. Cancer Discov. 9, 176–198. 10.1158/2159-8290.cd-18-1177.

31. Sun, H., Zou, J., Chen, L., Zu, X., Wen, G., and Zhong, J. (2017). Triple-negative breast cancer and its association with obesity. Mol. Clin. Oncol. 7, 935–942. 10.3892/mco.2017.1429.

32. van den Brandt, P.A., Spiegelman, D., Yaun, S.-S., Adami, H.-O., Beeson, L., Folsom, A.R., Fraser, G., Goldbohm, R.A., Graham, S., Kushi, L., et al. (2000). Pooled Analysis of Prospective Cohort Studies on Height, Weight, and Breast Cancer Risk. Am. J. Epidemiology 152, 514–527. 10.1093/aje/152.6.514.

33. Hopkins, B.D., Goncalves, M.D., and Cantley, L.C. (2016). Obesity and Cancer Mechanisms: Cancer Metabolism. J. Clin. Oncol. 34, 4277–4283. 10.1200/jco.2016.67.9712.

34. Wu, L., Wu, D., Ning, J., Liu, W., and Zhang, D. (2019). Changes of N6-methyladenosine modulators promote breast cancer progression. BMC Cancer 19, 326. 10.1186/s12885-019-5538-z.

35. Wang, H., Xu, B., and Shi, J. (2020). N6-methyladenosine METTL3 promotes the breast cancer progression via targeting Bcl-2. Gene 722, 144076. 10.1016/j.gene.2019.144076.

36. Huang, H., Weng, H., Deng, X., and Chen, J. (2019). RNA Modifications in Cancer: Functions, Mechanisms, and Therapeutic Implications. Annu. Rev. Cancer Biol. 4, 1–20. 10.1146/annurev-cancerbio-030419-033357.

37. Fan, Y., Lv, X., Chen, Z., Peng, Y., and Zhang, M. (2023). m6A methylation: Critical roles in aging and neurological diseases. Front. Mol. Neurosci. 16, 1102147. 10.3389/fnmol.2023.1102147.

38. Huang, Y., Xue, Q., Chang, J., Wang, Y., Cheng, C., Xu, S., Wang, X., and Miao, C. (2023). M6A methylation modification in autoimmune diseases, a promising treatment strategy based on epigenetics. Arthritis Res. Ther. 25, 189. 10.1186/s13075-023-03149-w.

39. Mao, Y., Dong, L., Liu, X.-M., Guo, J., Ma, H., Shen, B., and Qian, S.-B. (2019). m6A in mRNA coding regions promotes translation via the RNA helicase-containing YTHDC2. Nat. Commun. 10, 5332. 10.1038/s41467-019-13317-9.

40. Liu, Y.-L., Chou, C.-K., Kim, M., Vasisht, R., Kuo, Y.-A., Ang, P., Liu, C., Perillo, E.P., Chen, Y.-A., Blocher, K., et al. (2019). Assessing metastatic potential of breast cancer cells based on EGFR dynamics. Sci. Rep. 9, 3395. 10.1038/s41598-018-37625-0.

41. Zhang, R.D., Fidler, I.J., and Price, J.E. (1991). Relative malignant potential of human breast carcinoma cell lines established from pleural effusions and a brain metastasis. Invasion metastasis 11, 204–215.

42. Wang, Y., Li, Y., Toth, J.I., Petroski, M.D., Zhang, Z., and Zhao, J.C. (2014). N6-methyladenosine modification destabilizes developmental regulators in embryonic stem cells. Nat. Cell Biol. 16, 191–198. 10.1038/ncb2902.

43. Yang, Y., Fan, X., Mao, M., Song, X., Wu, P., Zhang, Y., Jin, Y., Yang, Y., Chen, L.-L., Wang, Y., et al. (2017). Extensive translation of circular RNAs driven by N6-methyladenosine. Cell Res. 27, 626–641. 10.1038/cr.2017.31.

44. Zeng, Z., Pan, Q., Sun, Y., Huang, H., Chen, X., Chen, T., He, B., Ye, H., Zhu, S., Pu, K., et al. (2023). METTL3 protects METTL14 from STUB1 mediated degradation to maintain m6A homeostasis. EMBO Rep. 24, e55762. 10.15252/embr.202255762.

45. Villa, E., Sahu, U., O’Hara, B.P., Ali, E.S., Helmin, K.A., Asara, J.M., Gao, P., Singer, B.D., and Ben-Sahra, I. (2021). mTORC1 stimulates cell growth through SAM synthesis and m6A mRNA-dependent control of protein synthesis. Mol. Cell 81, 2076–2093.e9. 10.1016/j.molcel.2021.03.009.

46. Armstrong, D.K., Kaufmann, S.H., Ottaviano, Y.L., Furuya, Y., Buckley, J.A., Isaacs, J.T., and Davidson, N.E. (1994). Epidermal growth factor-mediated apoptosis of MDA-MB-468 human breast cancer cells. Cancer Res. 54, 5280–5283.

47. Hendra, C., Pratanwanich, P.N., Wan, Y.K., Goh, W.S.S., Thiery, A., and Göke, J. (2022). Detection of m6A from direct RNA sequencing using a multiple instance learning framework. Nat. Methods 19, 1590–1598. 10.1038/s41592-022-01666-1.

48. Uzonyi, A., Dierks, D., Nir, R., Kwon, O.S., Toth, U., Barbosa, I., Burel, C., Brandis, A., Rossmanith, W., Hir, H.L., et al. (2023). Exclusion of m6A from splice-site proximal regions by the exon junction complex dictates m6A topologies and mRNA stability. Mol. Cell 83, 237–251.e7. 10.1016/j.molcel.2022.12.026.

49. Yang, X., Triboulet, R., Liu, Q., Sendinc, E., and Gregory, R.I. (2022). Exon junction complex shapes the m6A epitranscriptome. Nat. Commun. 13, 7904. 10.1038/s41467-022-35643-1.

50. He, P.C., Wei, J., Dou, X., Harada, B.T., Zhang, Z., Ge, R., Liu, C., Zhang, L.-S., Yu, X., Wang, S., et al. (2023). Exon architecture controls mRNA m6A suppression and gene expression. Science 379, 677–682. 10.1126/science.abj9090.

51. Yan, C., Xiong, J., Zhou, Z., Li, Q., Gao, C., Zhang, M., Yu, L., Li, J., Hu, M.-M., Zhang, C.-S., et al. (2023). A cleaved METTL3 potentiates the METTL3–WTAP interaction and breast cancer progression. eLife 12, RP87283. 10.7554/elife.87283.

52. Philippe, L., van den Elzen, A.M.G., Watson, M.J., and Thoreen, C.C. (2020). Global analysis of LARP1 translation targets reveals tunable and dynamic features of 5′ TOP motifs. Proc. Natl. Acad. Sci. 117, 5319–5328. 10.1073/pnas.1912864117.

53. Cockman, E., Anderson, P., and Ivanov, P. (2020). TOP mRNPs: Molecular Mechanisms and Principles of Regulation. Biomolecules 10, 969. 10.3390/biom10070969.

54. Schindelin, J., Arganda-Carreras, I., Frise, E., Kaynig, V., Longair, M., Pietzsch, T., Preibisch, S., Rueden, C., Saalfeld, S., Schmid, B., et al. (2012). Fiji: an open-source platform for biological-image analysis. Nat. Methods 9, 676–682. 10.1038/nmeth.2019.

