## Supplemental data for "Fine-tuning m6A and METTL3 levels have profound impact on cellular proliferation and protein synthesis"

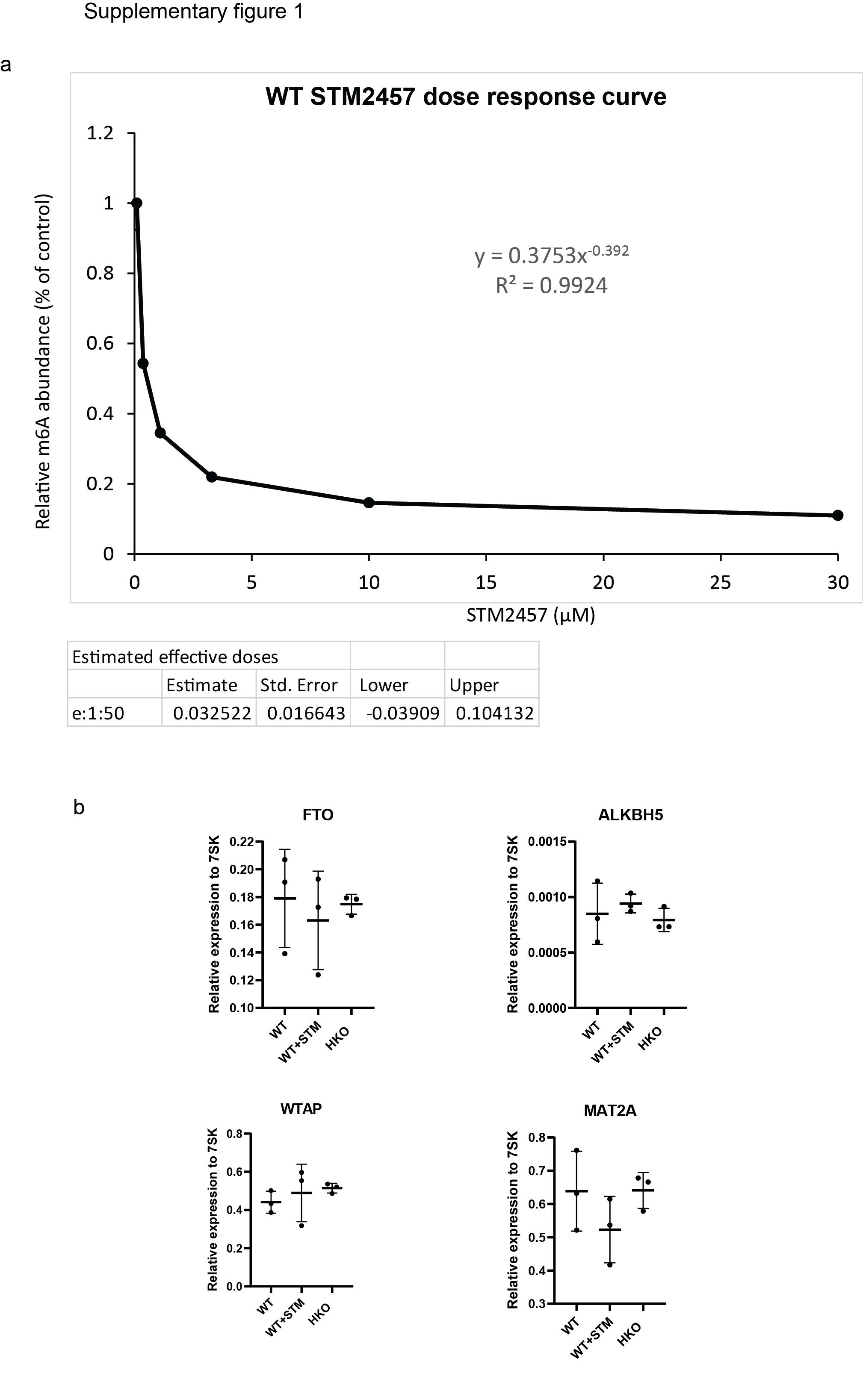

**Supplementary figure 1**

**a)** Dose response of STM2457 inhibition of METTL3 activity on MDA-MB-468 cells after 24h treatment showing relative m6A/A ratio in polyadenylated RNA as measured by LC–MS/MS (mean, n=3) Estimated effective dose e1:50, half maximum inhibitory concentration. b) Quantification of FTO, ALKBH5, WTAP and MAT2A mRNA levels in WT, WT treated with 0.145 µM STM2457 (WT+STM) and HKO cells by qRT-PCR. Data show the average (n=3) ± sem relative to RN7SK (7SK). P-value calculated by one-way ANOVA followed by Tukey HSD multiple comparison test.

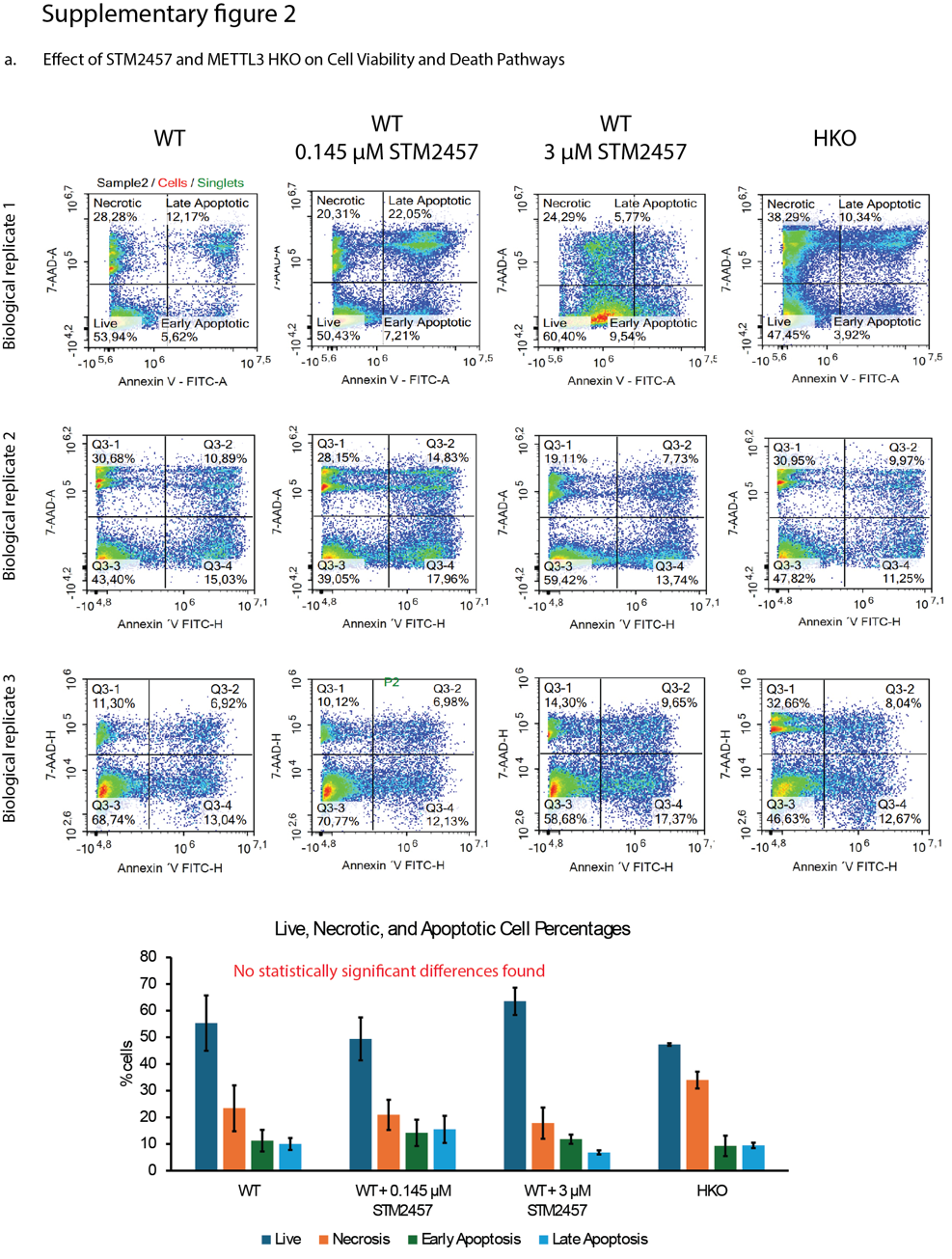

**Supplementary Figure 2** Partial knockout or inhibition of METTL3 does not significantly affect cell viability or cell death in MDA-MB-468 cells.

(A) Flow cytometry plots of MDA-MB-468 cells stained with Annexin V-FITC and 7-AAD, showing live, necrotic, early apoptotic, and late apoptotic populations across three biological replicates. Conditions include untreated wild-type (WT), WT treated with 0.145 µM STM2457, WT treated with 3 µM STM2457, and a CRISPR-generated heterozygous METTL3 knockout (HKO). (B) Quantification of cell populations from panel A. Data represent mean percentages across replicates. Statistical analysis using two-way ANOVA followed by Tukey’s HSD test revealed no significant differences between conditions, indicating that partial METTL3 inhibition or knockout does not impact cell viability or death.

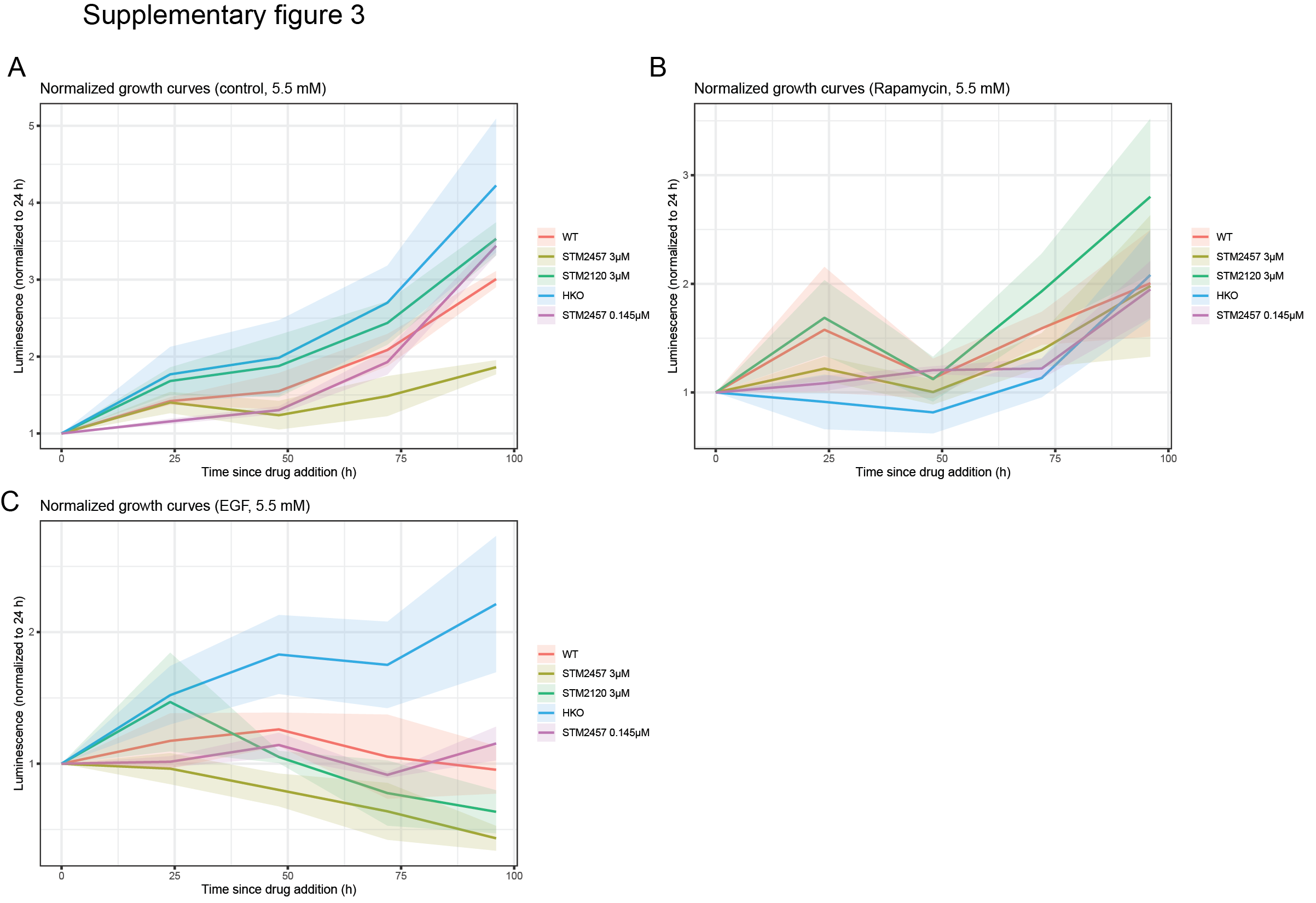
**Supplementary Figure 3. Normalised growth dynamics of MDA-MB-468 cells.**

Growth trajectories were normalized to the 24 h time point (24 h = 1; subsequent time points normalised to baseline) under (A) untreated, (B) EGF, and (C) rapamycin conditions, all at 5.5 mM glucose. Normalised curves highlight dynamic differences underlying the absolute AUC measurements while minimizing variability due to seeding density and plate effects. Normalized AUC values and associated statistics are provided in the Supplementary Data.

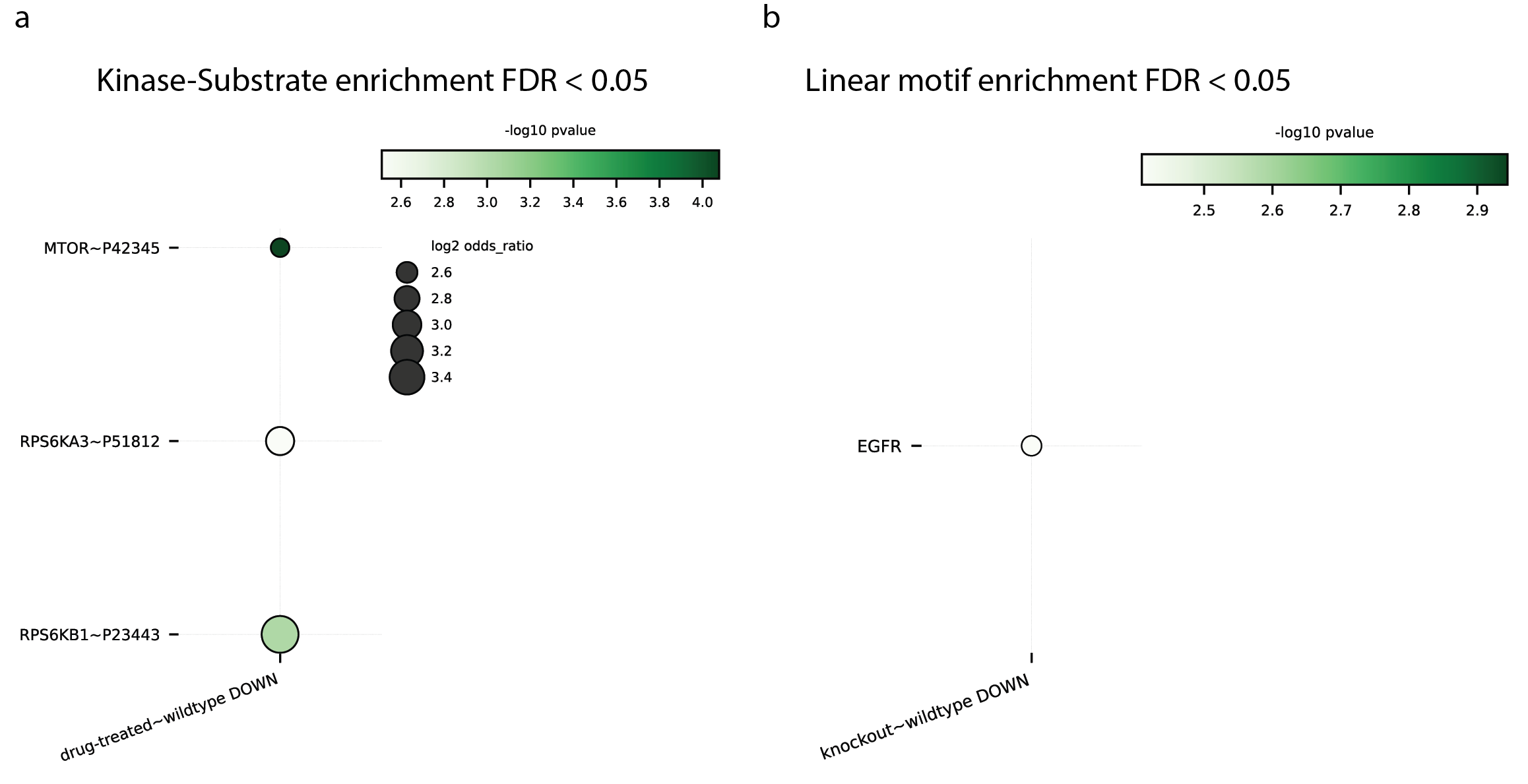

**Supplementary Figure 4 Protein Phosphorylation ontology. (a–b)** Kinase-substrate (a) and linear motif (b) enrichment of proteins undergoing differential phosphorylation upon partial reduction of METTL3 abundance or activity. Enrichment was performed using Mixed Permutation analysis (FDR < 0.05). Only conditions with more than one significantly enriched term (adjusted p-value < 0.05, ≥2 proteins per kinase or motif) are shown. Conditions without significant terms are omitted.

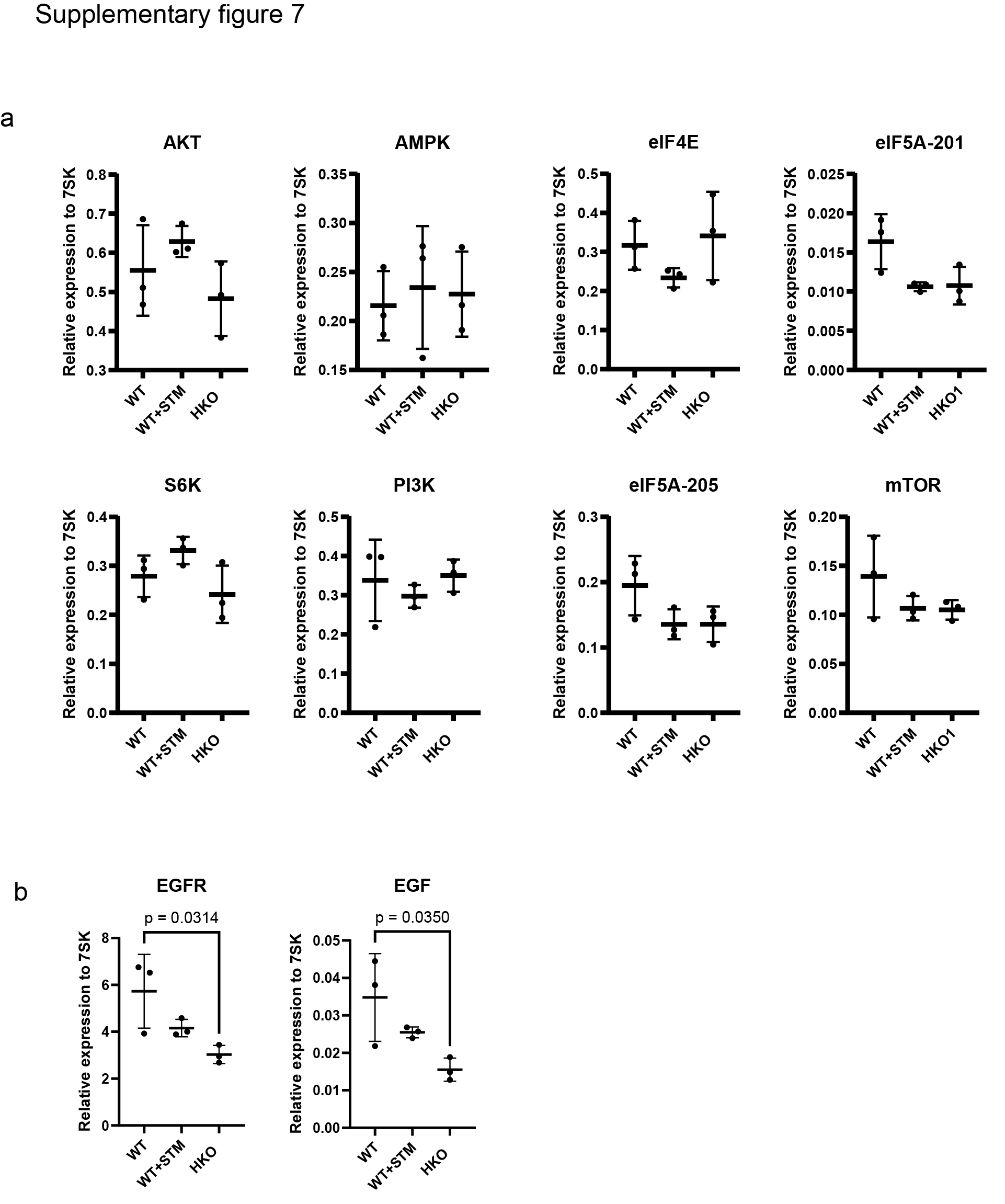
**Supplementary Figure 5**

**a)** Quantification of the mTOR signalling pathway components PI3K, AKT, mTOR, eIF4E, S6K as well as the energy sensor, AMPK, the two splice isoforms of the translation factor eIF5A-201 and eIF-205 mRNA levels by qRT-PCR, comparing samples with or without METTL3 inhibition/knockout. Data represent the average (n=3) ± SEM, normalized to RN7SK (7SK). P-values were calculated using one-way ANOVA followed by Tukey HSD multiple comparison test. **b)** Q-RT PCR quantification of the EGFR and its signalling molecule EGF. Data represent the average (n=3) ± SEM, normalized to RN7SK (7SK). P-values were calculated using one-way ANOVA followed by Tukey HSD multiple comparison test.

**Supplementary Table 1**

Significant differentially methylated kmers sites by xPore from Nanopore Direct RNA sequencing of WT MDA-MB-468 cells and METTL3 heterozygous knockout cells (HKO) (P-value <0.05, Z-score >1.9, kmers filtered for DRACH motifs only):

**Supplementary Table 2**

Significant differentially methylated sites identified by Nanocompore from Nanopore Direct RNA sequencing of WT MDA-MB-468 cells and METTL3 heterozygous knockout cells (HKO) and using the following statistical tests on current intensity and with a sequence context of 2, GMM logit, Kolmogorov Smirnov, Mann Whitney, and T-test (P-value <0.01):

**Supplementary Table 3**

m6Anet predicted m6A methylated sites from Direct RNA sequencing of WT MDA-MB-468 cells and METTL3 heterozygous knockout cells (HKO)

**Supplementary Table 4**

Gene ontology analysis of Nanocompore GMM logit context 2, KS current intensity context2, and xPore significant differentially modified genes. Includes both GO Terms and KEGG/Reactome terms.

**Supplementary Table 5**

Oligonucleotides used in this study
